# CYK-1/Formin activation in cortical RhoA signaling centers promotes organismal left-right symmetry breaking

**DOI:** 10.1101/2021.01.08.425924

**Authors:** Teije C. Middelkoop, Júlia Garcia-Baucells, Porfirio Quintero-Cadena, Lokesh G. Pimpale, Shahrzad Yazdi, Paul Sternberg, Peter Gross, Stephan W. Grill

**Affiliations:** Max Planck Institute of Molecular Cell Biology and Genetics, Dresden, Germany; Biotechnology Center, TU Dresden, Dresden, Germany; Cluster of Excellence Physics of Life, TU Dresden, Germany; California Institute of Technology, Division of Biology and Biological Engineering, Pasadena, USA; Massachusetts Institute of Technology, Department of Materials Science and Engineering, Cambridge, USA

## Abstract

1

Proper left-right symmetry breaking is essential for animal development and in many species the actin cytoskeleton plays an instrumental role in this process. Active torque generation in the actomyosin layer promotes left-right symmetry breaking in *C. elegans* embryos by driving chiral counter-rotating cortical flows. While both Formins and Myosins have been implied in left-right symmetry breaking, and both can rotate actin filaments *in vitro*, it remains unclear if active torques in the actomyosin cortex are generated by Formins, Myosins, or both. We combined the strength of *C. elegans* genetics with quantitative imaging and thin film, chiral active fluid theory to show that, while Non-Muscle Myosin II activity drives cortical actomyosin flows, it is permissive for chiral counter-rotation and dispensable for chiral symmetry breaking of cortical flows. Instead, we find that CYK-1/Formin activation in RhoA foci is instructive for chiral counter-rotation and promotes in-plane, active torque generation in the actomyosin cortex. Notably, we observe that artificially generated large active RhoA patches undergo rotations with consistent handedness in a CYK-1/Formin-dependent manner. Altogether, we conclude that, CYK-1/Formin-dependent active torque generation facilitates chiral symmetry breaking of actomyosin flows and drives organismal left-right symmetry breaking in the nematode worm.

**Significance:** Active torque generation in the actin cytoskeleton has been implicated in driving left-right symmetry breaking of developing embryos, but which molecules generate the active torque and how active torque generation is organized subcellularly remains unclear. This study shows that cortical Formin, recruited to cortical regions where RhoA signaling is active, promotes active torque generation in the actomyosin layer. We find that active torque tends to locally rotate the cortex in a clockwise fashion, which drives the emergence of chiral counter-rotating flows with consistent handedness and facilitates left-right symmetry breaking of *C. elegans* embryos.

## 3 Introduction

The emergence of left-right asymmetry is essential for normal animal development and in the vast majority of animal species, one type of handedness is dominant [1]. The actin cytoskeleton plays an instrumental role in establishing the left-right asymmetric body plan of invertebrates like fruit flies [2, 3, 4, 5, 6], nematodes [7, 8, 9, 10, 11, 12] and pond snails [13, 14, 15]. Moreover, an increasing number of studies now demonstrate that vertebrate left-right patterning also depends on a functional actomyosin cytoskeleton [14, 16, 17, 18, 19, 20, 21, 22]. Actomyosin-dependent chiral behaviour has even been reported in isolated cells [23, 24, 25, 26, 27, 28, 29] and such cell-intrinsic chirality has been shown to promote left-right asymmetric morphogenesis of tissues [30, 31], organs [21, 32] and entire embryonic body plans [13, 14, 33, 34]. Active force generation in the actin cytoskeleton is responsible for shaping cells and tissues during embryo morphogenesis. Torques are rotational forces with a given handedness and it has been proposed that in plane, active torque generation in the actin cytoskeleton drives chiral morphogenesis [7, 8, 35, 36].

What could be the molecular origin of these active torques? The actomyosin cytoskeleton consists of actin filaments, actin-binding proteins and Myosin motors. Actin filaments are polar polymers with a right-handed helical pitch and are therefore chiral themselves [37, 38]. Due to the right-handed pitch of filamentous actin (F-actin) both processive Myosins, like Myosin V, and non-processive myosins, like Myosin II, can rotate actin filaments along their long axis while pulling on them [34, 39, 40, 41, 42, 43]. Similarly, when physically constrained, members of the Formin family rotate actin filaments along their long axis while elongating them [44, 45]. In both cases the handedness of this rotation is determined by the helical nature of the actin polymer, and is therefore invariant. From this it follows that both Formins and Myosins are a potential source of molecular torque generation that could drive cellular and organismal chirality. Indeed chiral processes across different length scales, and across species, are dependent on Myosins [19], Formins [14, 15, 27, 46], or both [7, 8, 21, 47]. For chiral processes to emerge, active torque generation must be aligned with an axis of broken symmetry [1, 35]. It is however unclear how Formins and Myosins contribute to active torque generation and the emergence chiral processes in developing embryos.

In our previous work we showed that the actomyosin cortex of specific embryonic blastomeres undergoes chiral counter-rotations with consistent handedness [7, 36]. These chiral actomyosin flows can be faithfully recapitulated using active chiral fluid theory that describes the actomyosin layer as a thin film, active gel that generates active torques [7, 48, 49]. Chiral counter-rotating cortical flows reorient the cell division axis during cytokinesis, which is essential for normal left-right symmetry breaking [7, 50, 51]. Moreover, cortical counter-rotations with the same handedness have been observed in Xenopus one-cell embryos [33] suggesting that chiral counter-rotations are conserved among distant species. Chiral counter-rotating actomyosin flow in *C. elegans* blastomeres is driven by RhoA signaling and is dependent on Non-Muscle Myosin II motor proteins [7]. Moreover, the Formin CYK-1 has been implicated in actomyosin flow chirality during early polarization of the zygote as well as during the first cytokinesis [52, 53]. However, despite of having identified a role for Myosins and Formins, both of which could generate torque at the molecular level, the underlying mechanism by which active torques are generated remains elusive.

Here we show that the diaphanous-like Formin, CYK-1/Formin, is a critical determinant for the emergence of actomyosin flow chirality, while Non-Muscle Myosin II (NMY-2) plays a permissive role. Our results show that cortical CYK-1/Formin is recruited to active RhoA signaling foci and promotes active torque generation, which in turn tends to locally rotate the actomyosin cortex clockwise. In the highly connected actomyosin meshwork, a gradient of these active torques drives the emergence of chiral counter-rotating cortical flows with uniform handedness, which are essential for proper left-right symmetry breaking. Together, these results provide mechanistic insight into how Formin-dependent torque generation drives cellular and organismal left-right symmetry breaking.

## 4 Results

### CYK-1/Formin is a critical determinant of actomyosin flow chirality

Members of the Formin family have been implicated in chiral processes in multiple developmental contexts [14, 15, 21, 46, 47]. Moreover, the *C. elegans* Diaphanous-like Formin CYK-1/Formin was picked up in a candidate screen for chiral cortical flow phenotypes [52]. Therefore, we first asked whether the chiral counter-rotating actomyosin flows observed during anteroposterior polarisation of the *C. elegans* zygote are dependent on CYK-1/Formin. In order to perturb CYK-1/Formin protein function we made use of a temperature sensitive *cyk-1/Formin* loss of function mutant which yields non-functional CYK-1/Formin protein at the restrictive temperature of 25°C [54]. Polarizing actomyosin flows were recorded at the restrictive temperature in controls and *cyk-1/Formin* mutants by imaging the cortical surface of one-cell embryos producing endogenously-tagged Non-Muscle Myosin II (NMY-2::GFP) [55]. Subsequently, cortical flows were quantified using particle image velocimetry (PIV) and a mean velocity profile was obtained by averaging velocity vectors over a time period of 100 seconds during polarizing flow. Although cortical flow in control embryos is mainly directed along the anteroposterior (AP) axis, we find that the anterior and posterior cortical halves counter-rotate relative to each other (Movie 1, Figure 1A-C). Notably, the handedness of the counter-rotation is consistent among embryos (Figure 1A-C) and these findings are in line with earlier results [7].

**Figure 1:**
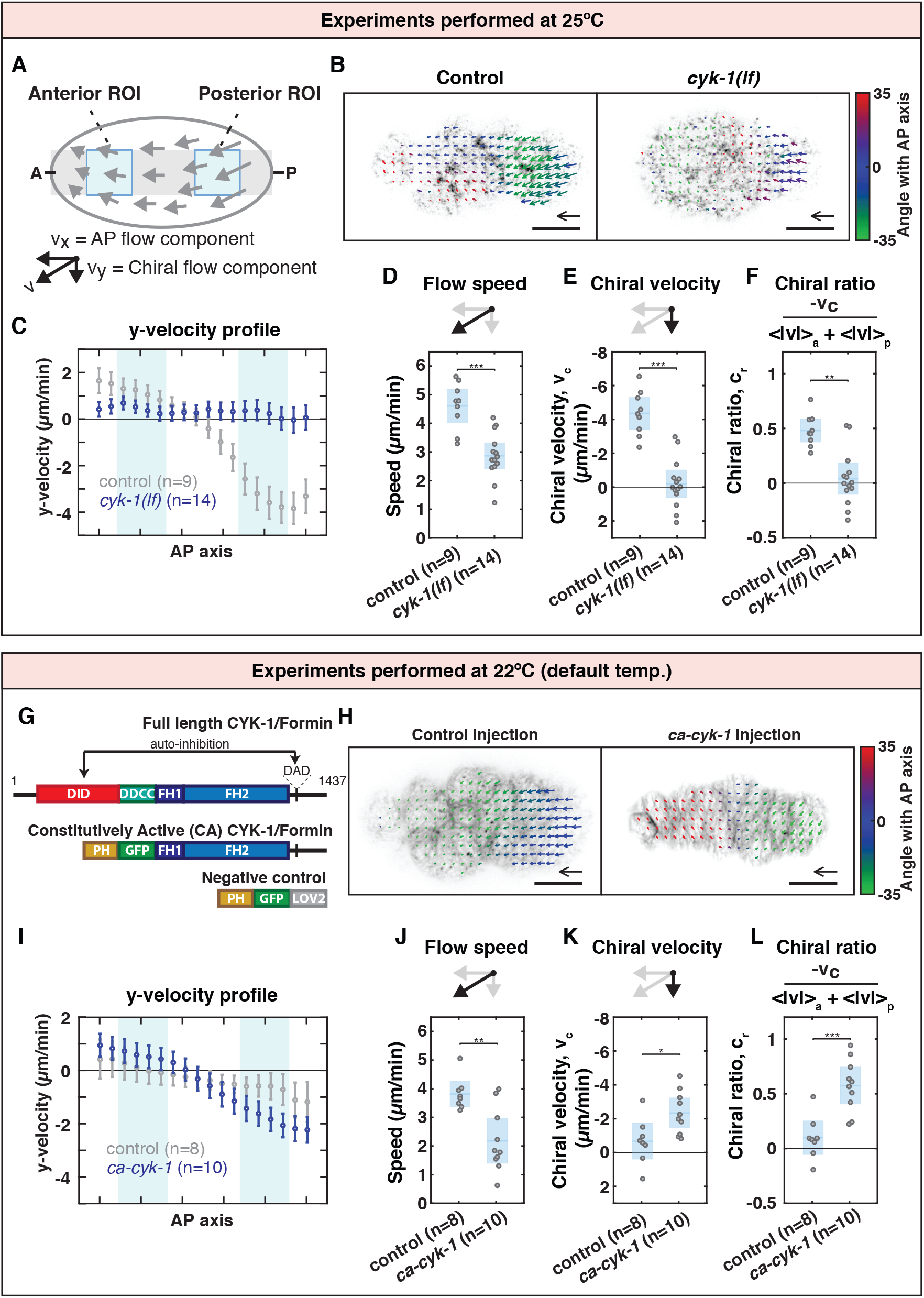
CYK-1/Formin is a determinant of actomyosin flow chirality. (**A**) Schematic of a *C. elegans* zygote during anteroposterior polarization. The grey stripe along the anteroposterior axis depicts the region used to obtain the flow velocity profiles. Blue boxes indicate the regions of interest that were used to calculate the mean flow speed, chiral velocity and chiral ratio. Results shown in (**B**) - (**F**) were obtained by imaging embryos at (25°C). (**B**) Time-averaged flow field overlaid on a fluorescent still image of NMY-2::GFP imaged at the cortical surface of a control and a *cyk-1(lf)* mutant embryo. Mean velocity vectors are color coded for their angle with the anteroposterior axis. Scale bar, 10 *µ*m. Velocity scale arrow, 20 *µ*m/min. (**C**) Mean y-velocity in 18 bins along the anteroposterior axis measured in a stripe as indicated in (**A**) averaged over embryos for control (grey) and for *cyk-1(lf)* (blue). Light blue areas mark bins 3-6 and 13-16 corresponding to the anterior and posterior ROIs respectively. Error bars, SEM. (**D**) Mean speed per embryo as defined as (*<* |v| >_a_ + *<* |v| >_p_)*/*2, where *<* |v| >_a_ and *<* |v| >_p_ are the spatial averages in the anterior and posterior regions of interest respectively. (**E**) Mean chiral velocity, v_c_, per embryo as defined as *<* v_y_ >_p_ − *<* v_y_ >_a_, where *<* v_y_ >_a_ and *<* v_y_ >_p_ are the spatially averaged y-velocities in the anterior and posterior regions of interest respectively. (**F**) Mean chiral ratio per embryo, defined as 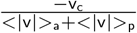 (**G**) Domain overview of CYK-1/Formin/mDia (upper schematic) and the recombinant constructs generated in this study (CA-CYK-1/Formin, middle and PH-GFP-LOV2, lower schematic). DID=Dia Inhibitory Domain, DD=Dimerization Domain, CC=Coiled Coil, FH=Formin Homology, DAD=Diaphanous Autoregulatory Domain, PH=Pleckstrin Homology, GFP=Green Fluorescent Protein, LOV2=Light-Oxygen-Voltage 2 domain. Results shown in (**H**) - (**L**) were obtained by imaging embryos at (22°C). (**H**) Time-averaged flow field overlaid on a fluorescent still image of Lifeact-mKate2 imaged at the cortical surface of an embryo derived from control (*ph-gfp-lov2*) injection and from (*ca-cyk-1/Formin*) injection. Vector color coding, scale bar and vector scale arrow as in (**B**). (**I**) Mean y-velocity profile in 18 bins along the anteroposterior axis, in embryos derived from control injection (grey) and from *ca-cyk-1/Formin* injection (blue). Error bars, SEM. Mean flow speed (**J**), chiral velocity (**K**) and chiral ratio (**J**) in embryos derived from control injection and from *ca-cyk-1/Formin* injection. Blue in (**D**)-(**F**) and **J**)-(**L** depicts the mean over embryos at 95% confidence. Significance testing: * represents p*<*=0.05, ** represents p*<*=0.01, *** represents p*<*=0.001 (Wilcoxon rank sum test). n indicates the number of embryos.

To quantify chiral counter-rotation of the flow we first decomposed velocity vectors into a component along the AP axis (x-velocity, v_x_) and a chiral component perpendicular to the AP axis (y-velocity, v_y_)(Figure 1A). Subsequently, we computed the chiral velocity, denoted v_c_, by subtracting the averaged y-velocity in the anterior from the averaged y-velocity in the posterior (v_c_ =*<* v_y_ >_p_ − *<* v_y_ >_a_)[7]. Non-perturbed control embryos display a v_c_ of −4.36 ± 1.00 *µ*m/min (mean ± 95% C.I. throughout the manuscript)) (Figure 1E). In contrast, *cyk-1/Formin* loss of function resulted in a v_c_ of −0.20 ± 0.82 *µ*m/min, indicating that the chiral counter-rotation was lost entirely (Movie2, Figure 1B, C, E). Note that the average flow speed (Figure 1D), x-velocity (Supplementary figure 1A-B) and flow coherence (Supplementary figure 1C) were significantly reduced in *cyk-1/Formin* mutants suggesting that, in addition to flow chirality, CYK-1/Formin affects multiple aspects of flow dynamics. In order to exclude that the reduced chiral velocity is simply due to a reduction in overall flow speed, we normalized the chiral velocity by the flow speed and define a chiral ratio, c, according to 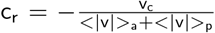, where *<* |v| >_a_ and *<*|v|>_p_ are the mean flow speed in the anterior and posterior, respectively. Note that the chiral ratio is positive with wild type flow handedness, and unperturbed control embryos display a chiral ratio of 0.48 ± 0.11 (Figure 1F). In contrast, the chiral ratio in *cyk-1/Formin* mutants is 0.04 ± 0.14 (Figure 1F) indicating that CYK-1/Formin is indeed required to generate actomyosin flows with a chiral counter-rotating component.

The flow behaviour in the control reported in Figure 1B-E is overall similar to our previous observations [7]. However, the chiral velocity, here measured at 25 °C and in a different genetic background, differs (−2.9 ± 0.3 *µ*m/min in [7] as opposed to −4.36 ± 1.00 *µ*m/min here, Figure 1C, E). We found that both the temperature and the genetic background affect the chiral velocity (see supplementary note and Supplementary figure 2). Throughout the remainder of this study we used the default temperature of 22°C in all experiments, unless explicitly stated otherwise.

We next asked whether CYK-1/Formin activity is sufficient to promote chiral counter-rotating flows in *C. elegans* zygotes. To this end, we sought to experimentally increase CYK-1/Formin activity. The overall domain structure of CYK-1/Formin is similar to other Diaphanous-like formins (Figure 1G)[56], where the Formin Homology domains (FH1 and FH2) at the C-terminus display the enzymatic activity required for actin polymerization and nucleation [57, 58]. The FH domains are flanked by a C-terminal DAD peptide and an N-terminal DID domain that associate intramolecularly, rendering the protein inactive. Formin activators like Rho GTPases release this auto-inhibition by binding to the DID and a neighbouring GTPase binding region [59, 60, 61, 62, 63]. It has been previously reported that deletion of the DID domain of Diaphanous-related Formins, results in constitutively active protein *in vivo* [64, 65]. Therefore, in order to experimentally increase cortical CYK-1/Formin activity in *C. elegans* embryos, we generated an N-terminally truncated CYK-1/Formin construct (residues 700-1437), deleting everything upstream of the FH1 domain, and replaced the N-terminus with a GFP-tagged Pleckstrin Homology (PH-GFP) domain that facilitates membrane localization (Figure 1G). We will refer to this construct as Constitutively Active CYK-1/Formin (CA-CYK-1/Formin).

We expressed *ca-cyk-1/Formin* transiently (see methods) in the adult gonad of worms carrying a *lifeact::mKate2* transgene that are otherwise wild type. This allowed us to obtain zygotes with variable levels of CA-CYK-1/Formin. Introducing high levels of CA-CYK-1/Formin in embryos expressing Lifeact-mKate2 (a marker for F-actin) revealed numerous cortical anomalies. In many embryos cortical actomyosin flow direction changed repeatedly and this was accompanied by repeated reorientations of local actin filament alignment (Movie 4, see also the supplementary note). Such defects were never observed in embryos derived from negative control injections (*ph-gfp-lov2*, Movie 3). Moreover, expression of CA-CYK-1/Formin resulted in an increase in the cortex-to-cytoplasm ratio of Lifeact-mKate2, consistent with an elevated cortical abundance of F-actin (Supplementary figure 1D,G-I). Altogether, these findings demonstrate that the CA-CYK-1/Formin construct is indeed constitutively active.

We next sought to determine whether introducing CA-CYK-1/Formin affects actomyosin flow chirality. Because high levels of constitutively active CYK-1/Formin strongly perturb cortex physiology (Movie 4), we analyzed embryos producing low levels of CA-CYK-1/Formin. This is possible because we expressed *ca-cyk-1* transiently, and because cortical levels can be measured since CA-CYK-1/Formin is GFP-labeled (Supplementary figure 1D-E). Introduction of low levels of CA-CYK-1/Formin resulted in a significant increase in both the chiral counter-rotation of the flow and the chiral ratio (Movie 5, Figure 1H, I, K, L, Supplementary figure 1E-F), indicating that CA-CYK-1/Formin activity is sufficient to promote chiral counter-rotating flows. We note that the mean cortical flow speed, and the anteroposterior component of the flow (v_x_), were substantially reduced when compared to control injections (Movie 3, Movie 5, Figure 1J, Supplementary figure 1J-K), indicating that constitutive CYK-1 activity has additional effects on cortex dynamics. We also note that the chiral velocity and chiral ratio in the negative control (*ph-gfp-lov2* plasmid injected into *lifeact-mKate2* worms) are reduced when compared to those observed in the control imaged at 25 ° C (Figure 1B-F). This is consistent with both temperature and genetic background impacting on actomyosin flow chirality (see supplementary note and Supplementary figure 2). Together, given that the chiral ratio is increased upon introduction of CA-CYK-1/Formin, these results lend credence to the statement that CYK-1/Formin affects the strength of chiral counter-rotation. We conclude that CYK-1/Formin is a critical determinant for actomyosin flow chirality.

### CYK-1/Formin promotes torque generation in the actomyosin layer

Given that CYK-1/Formin determines actomyosin flow chirality, we hypothesized that CYK-1/Formin itself could generate active torques. Alternatively, given that strong loss or gain of CYK-1/Formin function results in a reduction of cortical flow speed (Figure 1D, J), CYK-1/Formin could play an indirect, permissive role in the emergence of chiral cortical flows, by generating the cortical F-actin needed to support active forces and torques. To discriminate between these two possibilities, we next performed weak perturbation *cyk-1(RNAi)* experiments, in which we vary the depletion strength by varying the RNAi feeding time (ranging from 6-24 hrs). Subsequently, we quantified the cortical flow dynamics upon mild deviations from wild type CYK-1 levels, equivalent to determining the linear response to a small perturbation. These *cyk-1(RNAi)* treatments were performed on embryos producing endogenously labeled NMY-2 (NMY-2::mKate2) to measure cortical flows, and CYK-1/Formin (CYK-1/Formin::GFP) to measure cortical levels of CYK-1/Formin (Figure 2A-C). We hypothesized that if CYK-1/Formin affects flow chirality indirectly by modulating cortex structure, both flow speed and flow chirality will decrease with decreasing cortical CYK-1/Formin levels. Alternatively, if CYK-1/Formin generates active torques itself, flow chirality, but not flow speed, will decrease with decreasing cortical CYK-1/Formin levels. Consistent with the latter hypothesis, we found that both the chiral counter-rotation velocity, v_c_ (Supplementary figure 3D), and the chiral ratio c_r_ of the flow (Figure 2D) correlated with cortical CYK-1/Formin levels (Spearman’s *ρ*=-0.44, p*<*0.0007 and *ρ*=0.54, p*<*0.00002, respectively), while we found no significant correlation between CYK-1/Formin levels and cortical flow speed (Spearman’s *ρ*=-0.23, p=0.08)(Figure 2E). We note that even a strong *cyk-1(RNAi)* perturbation did not affect the mean flow speed as did the *cyk-1/Formin* loss of function mutation described above. This apparent discrepancy is likely to be explained by residual CYK-1/Formin levels remaining, even upon *cyk-1(RNAi)* treatment. Together, our results show that, at physiological levels, CYK-1/Formin modulates flow chirality but not flow speed. These findings are consistent with a direct role for CYK-1/Formin in promoting active torque generation in the cortex.

**Figure 2:**
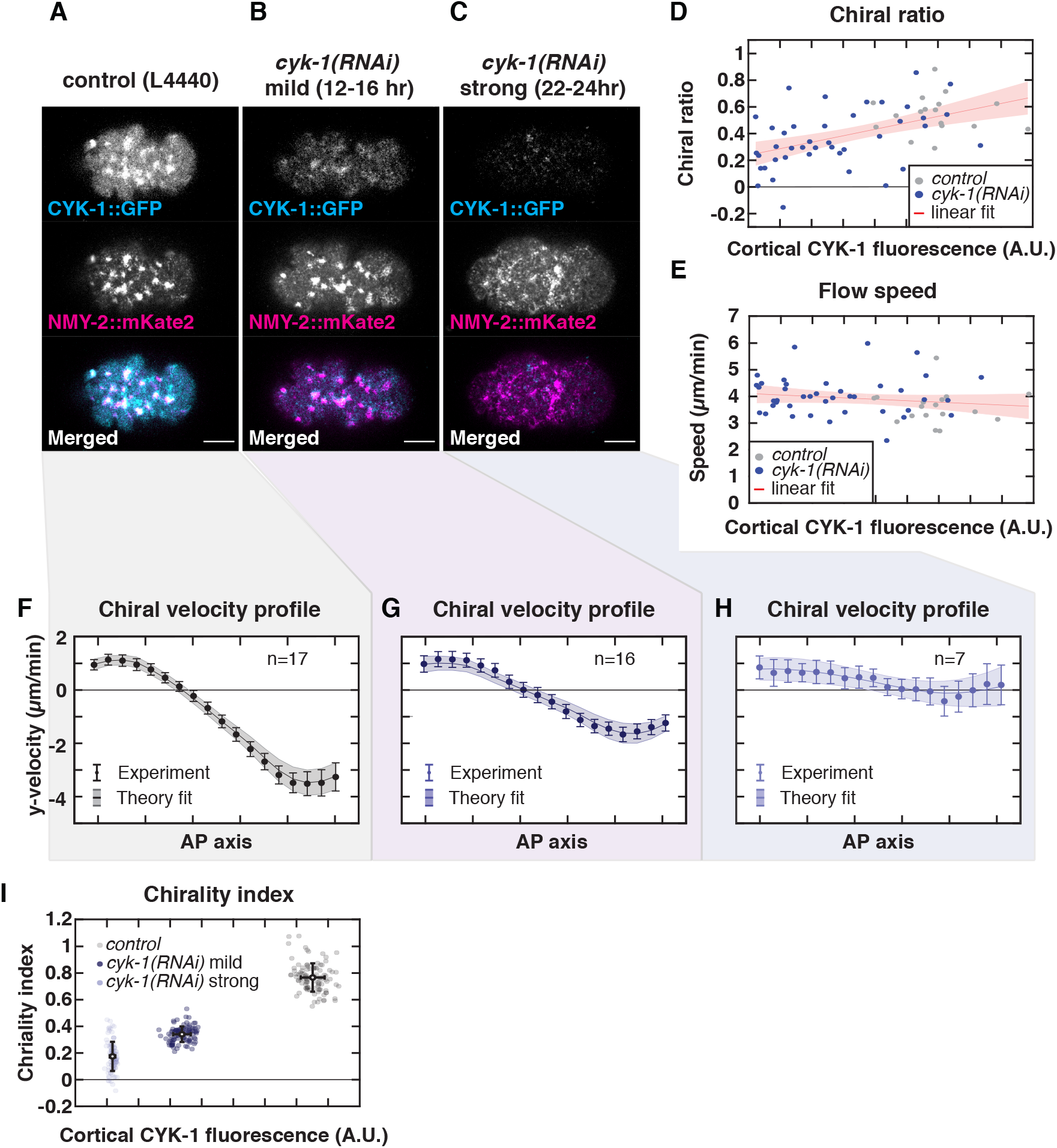
CYK-1/Formin promotes torque generation in the actomyosin cortex. (**A**)-(**C**) Representative cortical plane micrographs embryos producing CYK-1/Formin::GFP and NMY-2::mKate2 in (**A**) control (L4440), (**B**) mild *cyk-1(RNAi)* (12-16 hrs) and (**C**) strong *cyk-1(RNAi)* (22-24 hrs) during polarizing flows. Scale bar, 10 *µ*m. (**D**) Chiral ratio and (**E**) speed of the cortical flow plotted over the measured cortical CYK-1/Formin::GFP fluorescence in control (L4440, grey) and upon increasing strength of *cyk-1(RNAi)* (blue). Data points represent individual embryos. Red line with shaded region shows a linear fit with 95% confidence bounds. Chiral ratio, but not flow speed, correlates with cortical CYK-1/Formin::GFP (Spearman’s *ρ*=0.54, p*<*0.00002). control, n=17 embryos *cyk-1(RNAi)* n=41 embryos. (**F**)-(**H**) Mean y-velocity profile in 18 bins along the anteroposterior axis. Conditions as in (**A**)-(**C**). Circles with errorbars are the experimentally measured y-velocities averaged over embryos, with SEM. Solid line shows the mean y velocities derived from fitting the hydrodynamic model to 100 bootstrap samples with replacement. The shaded region displays the standard deviation of the mean, derived from bootstrapping. n indicates the number of embryos. (**I**) Chirality index plotted over the measured cortical CYK-1/Formin::GFP fluorescence in control (L4440, grey), mild *cyk-1(RNAi)* (12-16 hrs, dark blue) and strong *cyk-1(RNAi)* (22-24 hrs, light blue). Chirality index was obtained by fitting the hydrodynamic model to the mean of individual bootstrap samples with replacement. Simultaneously, in each bootstrap sample the mean cortical CYK-1/Formin::GFP fluorescence was calculated. Grey and blue data points represent individual bootstrap samples. Black points with errorbars display the mean over all bootstrap samples with standard deviation.

To determine whether CYK-1/Formin indeed promotes active torque generation, we next applied active chiral fluid theory [48, 49]. This coarse-grained theoretical description is built on the principle that the cortex tends to locally contract in response to active tension and tends to locally rotate clockwise (when viewed from the outside) in response to active torque generation. An anteroposterior gradient of active tension drives flows along the AP axis, while an anteroposterior gradient of active torques drives chiral-counter rotation of the flow. By fitting the experimentally measured flow profiles to the active chiral fluid model, the amount of active tension and active torque generation can be obtained. From this, a dimensionless measure for actomyosin flow chirality can be computed by taking the ratio of active torque density to active tension, denoted as the chirality index c_i_ [7]. Note that, although both the chirality index, c_i_ and the chiral ratio, c_r_ (introduced above) are measures for the relative strength of chirality, the chirality index is derived from fitting the theoretical model while the chiral ratio is measured directly from the obtained flow velocity profile. Fitting the theoretical model to the flow velocity profiles of control, mild and strong *cyk-1(RNAi)* yielded a good agreement between model and experiment (Figure 2F-H, Supplementary figure 3A-C). Moreover, we found that the chirality index c_i_ correlates with cortical CYK-1/Formin levels (Figure 2I). These results further substantiate our conclusion that CYK-1/Formin determines cortical flow chirality by promoting active torque generation in the actomyosin layer.

### CYK-1/Formin is required for organismal left-right symmetry breaking

Left-right symmetry breaking in *C. elegans* embryos occurs by a chiral skew of the ABa/ABp cell division axes in the 4-6 cell embryo [50, 51] and chiral counter-rotating actomyosin flows drive this event [7, 36]. Therefore, we next asked whether this cell division skew is dependent on *cyk-1/Formin*. Given that *cyk-1/Formin* is required for normal cytokinesis, *cyk-1/Formin* temperature sensitive mutants and controls were grown at permissive temperature (15°C) until the 4-cell stage and then shifted to the restrictive temperature (25°C) prior to ABa/ABp cytokinesis. In order to quantify cell division skews, we imaged embryos expressing a tubulin marker (mCherry::Tubulin) and used the orientation of the mitotic spindle as a proxy for the cell division axis (Figure 3A-B). Imaging the mitotic spindle during ABa/ABp cytokinesis revealed that the cell division skew in control embryos was 26.27 ± 1.74 and 23.62 ± 1.53 degrees for ABa and ABp, respectively (Movie 6, Figure 3B, D, E), which is consistent with previous reports [7, 36, 50, 51]. This spindle skew was strongly reduced in *cyk-1/Formin* mutants (7.60 ± 0.81 and 6.50 ± 1.05 degrees for ABa and ABp respectively, Movie 7, Figure 3C, D, E), demonstrating a role for *cyk-1/Formin* in left-right symmetry breaking. Given that chiral counter-rotating flows drive the spindle skew of ABa and ABp [7, 36], and given that we show here that CYK-1/Formin determines actomyosin flow chirality, we conclude that CYK-1/Formin facilitates left-right symmetry breaking by controlling actomyosin flow chirality.

**Figure 3:**
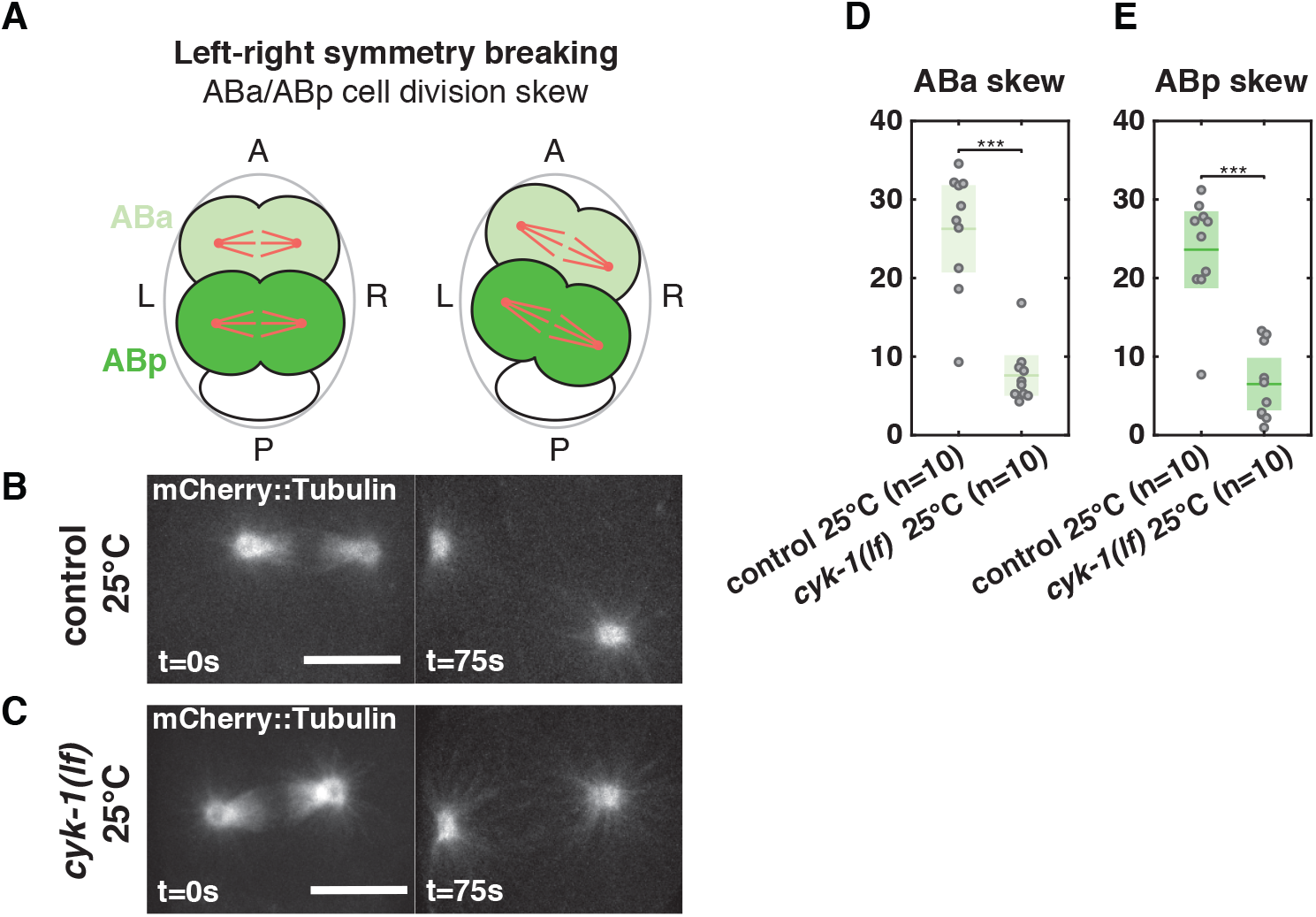
CYK-1/Formin is required for left-right symmetry breaking. (**A**) Schematic of a 4 cell embryo undergoing left-right symmetry breaking, viewed down the dorsoventral axis (anterior is up, left side is left). The ABa (light green) and ABp (dark green) blastomeres undergo a clockwise cell division skew such that their left and right daughter cells get displaced towards the anterior and posterior respectively. (**B**)-(**C**) Micrographs of the ABp spindle (marked with mCherry-Tubulin) at the beginning (t=0 sec, left) and at the end of anaphase B (t=75 sec, right) in a (**B**) control and a (**C**) temperature sensitive *cyk-1(lf)* mutant embryo. Imaging was performed at the restrictive temperature (25°C). Scale bar, 10 *µ*m. (**D**)-(**E**) Dotplots of the measured spindle skew angle of (**D**) the ABa and (**E**) the ABp cell. Spindle skew angle is defined as the difference between the spindle angle at the onset and at the end of anaphase B. Dots represent individual embryos, green shows the mean with the 95% confidence interval. Significance testing: *** represents p*<*=0.001 (Wilcoxon rank sum test). n indicates the number of embryos.

### CYK-1/Formin is a target of RhoA signaling

In many different contexts Formins act directly downstream of the RhoA GTPase [66]. Moreover, we showed previously that chiral actomyosin flows in the *C. elegans* embryo are controlled by RhoA signaling [7]. Therefore, we next asked whether CYK-1/Formin is a RhoA target in the *C. elegans* zygote. Analysis of endogenously labeled CYK-1/Formin (CYK-1/Formin::GFP) revealed that CYK-1/Formin localizes in distinct cortical foci that are depleted from the posterior during the cortical flow phase. This sets up a gradient of CYK-1/Formin::GFP along the anteroposterior axis (Movie 8, Figure 4A, B). The shape of this gradient, as well as the localization in foci, is similar to what is observed for other RhoA effectors (Movie 10, Supplementary figure 4C, D) [67, 68, 69, 70, 71]. We next tested whether the CYK-1/Formin foci are indeed regions of high RhoA activity. Earlier studies have used the AH-PH domain of ANI-1 fused to GFP as a marker for RhoA activity [71]. We generated an mCherry-tagged version of this probe and combined it with endogenously-tagged CYK-1/Formin (CYK-1/Formin::GFP). Dual color imaging during polarizing flow revealed a significant co-localization (Pearson *ρ*=0.35), indicating that CYK-1/Formin::GFP indeed localizes to regions of active RhoA (Movie 12, Figure 4C-D).

**Figure 4:**
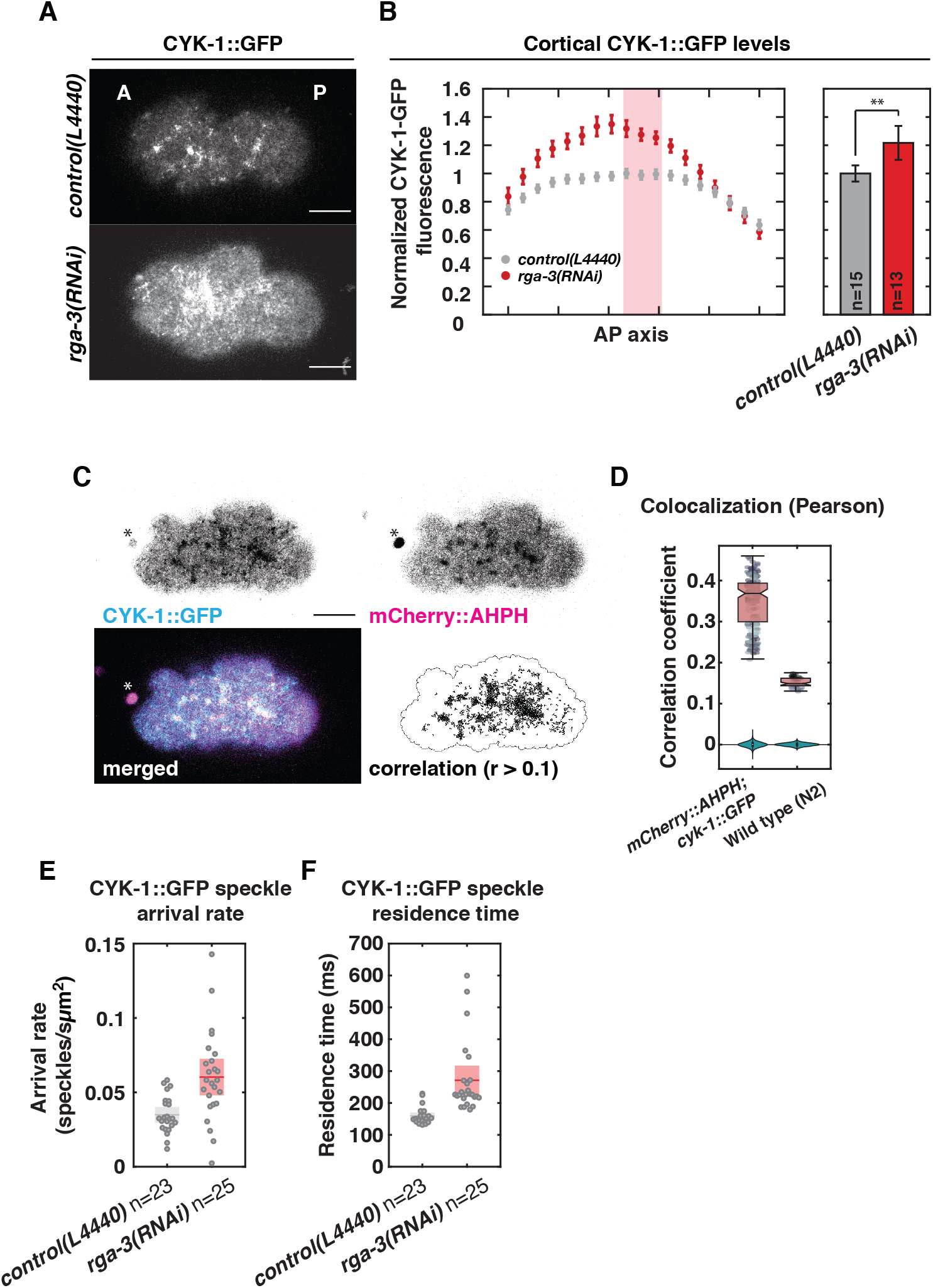
CYK-1/Formin is a RhoA target during polarizing flows. (**A**) Fluorescent micrographs of the cortical plane of embryos expressing CYK-1/Formin::GFP captured during polarizing flows in control (L4440, upper panel) and upon *rga-3(RNAi)* (lower panel). Scale bar, 10 *µ*m. (**B**) Left panel: Profile of the mean cortical CYK-1/Formin::GFP fluorescence in 18 bins along the anteroposterior axis in control (L4440, grey) and upon *rga-3(RNAi)*(red). Fluorescence levels were normalized to the mean levels in bins 9-11 (light red rectangle). Error bars, SEM. Right panel: bar diagram displaying the mean cortical CYK-1/Formin::GFP fluorescence measured in bins 9-11 (light red rectangle in the left panel). Error bars, 95% confidence interval. Significance testing: ** represents p *<*= 0.01 (Wilcoxon rank sum test). n indicates the number of embryos. (**C**) Micrographs (upper 2 panels and lower left) of the cortical plane of embryos producing CYK-1/Formin::GFP and mCherry::ANI-1(AHPH) captured during polarizing flow. Lower right panel displays regions of colocalization in black, that display a Pearson correlation coefficient larger than 0.1. Black line indicates the mask boundary and delineates the cortical surface that is in focus. Scale bar, 10 *µ*m. Asterix marks the polar body. (**D**) Pearson correlation coefficient computed for individual time frames (15 frames per embryo) in embryos producing CYK-1/Formin::GFP and mCherry::ANI-1(AHPH) (n=13 embryos) and non-transgenic wild type embryos to control for correlation of autofluorescence (n=4 embryos). Data points represent individual frames and a boxplot is overlaid (displaying the minimum, first quartile, median, third quartile and the maximum). As a negative control the correlation was computed using scrambled pixel values for one channel. This was done 100 times independently for both conditions. Violin plots display the distribution of these negative controls. (**E**) CYK-1/Formin::GFP speckle arrival rate and (**F**) residence time in control (L4440, grey, n=23 embryos) and upon *rga-3(RNAi)*(red). Data points represent individual embryos. Boxes indicate mean with 95% confidence interval.

If CYK-1/Formin is indeed a target of active RhoA, then cortical CYK-1/Formin levels will decrease with decreased RhoA signaling activity. To test this, we first reduced RhoA activity by RNAi of *ect-2*, which is the upstream activator of RhoA [72]. *ect-2(RNAi)* resulted in a subtle reduction of cortical CYK-1/Formin::GFP levels, and abolished its localization in cortical foci (Supplementary figure 4A-B). In addition, increasing overall RhoA activity by knocking down a negative regulator of RhoA signaling, *rga-3*, led to a substantial increase of cortical CYK-1/Formin levels and cortex-to-cytoplasm ratio (Movie 9, Figure 4A, B, Supplementary figure 4 E-G). Importantly, we observed a similar increase when analysing cortical levels of NMY-2::GFP (Movie 11, Supplementary figure 4C-D), a well known target of RhoA [72, 73, 74]. Altogether, the results presented here are consistent with CYK-1/Formin being a target of RhoA.

We next asked how active RhoA controls CYK-1/Formin localization dynamics at the molecular level. To this end we performed speckle microscopy [75, 76, 77] of endogenously labeled CYK-1/Formin::GFP in control embryos and upon *rga-3(RNAi)* treatment during polarizing actomyosin flows. To achieve sparse labeling of the fluorescent CYK-1/Formin::GFP pool we made use of the fact that the CYK-1/Formin::GFP signal photobleaches rapidly upon continuous imaging. After 50 second of imaging more than 98% of the fluorescent signal photobleached (Supplementary figure 5 A). Due to strong photobleaching, the remaining fluorescent CYK-1/Formin::GFP appeared as sparsely localized, dynamic speckles in the cortical plane. This allowed us to detect newly arriving cortical CYK-1/Formin molecules from which we measured the cortical arrival rate and the residence time.

To determine how active RhoA controls CYK-1/Formin dynamics, we tracked a total of 55612 CYK-1/Formin::GFP speckles in control (n=23 embryos) and 89080 in *rga-3(RNAi)* (n=25 embryos) and found that *rga-3(RNAi)* increases both arrival rate and residence time (Figure 4E-F). Importantly, the measured residence times in control (159 ± 11 ms) and *rga-3(RNAi)* (271 ± 46 ms) are much lower than the half life due to photobleaching (1246 ± 90 ms, Supplementary figure 5 B), indicating that photobleaching only results in a mild underestimation of the residence time. Our observations that active RhoA increases cortical CYK-1/Formin levels by increasing the arrival rate and residence time of CYK-1/Formin, are consistent with a recent study that showed that CYK-1/Formin acts downstream of RhoA pulses [78]. These results further substantiate our conclusion that CYK-1/Formin is RhoA target.

### RhoA controls actomyosin flow chirality via CYK-1/Formin activation

The results presented here show that CYK-1/Formin is a critical determinant of actomyosin flow chirality and that CYK-1/Formin is a RhoA target. In our earlier work we have demonstrated that chiral counter-rotating flows are driven by RhoA signaling and are dependent on the RhoA target Non-Muscle Myosin II (NMY-2) [7]. To further dissect the roles of NMY-2 and CYK-1/Formin in the emergence of chiral counter-rotating actomyosin flows, we asked which of these two RhoA targets is responsible for the hyperchirality phenotype associated with elevated RhoA activity. In order to experimentally increase RhoA signaling, we made use of a gain of function mutation in the upstream RhoA activator, *ect-2* [79]. While in wild type embryos the anteroposterior component of the flow is dominant and chiral counter-rotation is subtle, in *ect-2* gain of function mutants we observed the opposite: the anteroposterior component of the flow is strongly reduced and the chiral counter-rotation of the flow is dominant (Movie 13, Movie 14, Figure 5A, B, F, Supplementary figure 6A). These findings are similar to those observed upon strong *rga-3(RNAi)* (Movie 11). Surprisingly, despite increased RhoA signaling levels, the overall flow speed in *ect-2(gf)* mutants was only mildly increased when compared to wild type (Figure 5E). Hence, the increased RhoA activity in the *ect-2(gf)* mutant mainly acts to increase the chiral ratio (Figure 5G), e.g. the fraction of the total flow speed accounted for by the chiral velocity.

**Figure 5:**
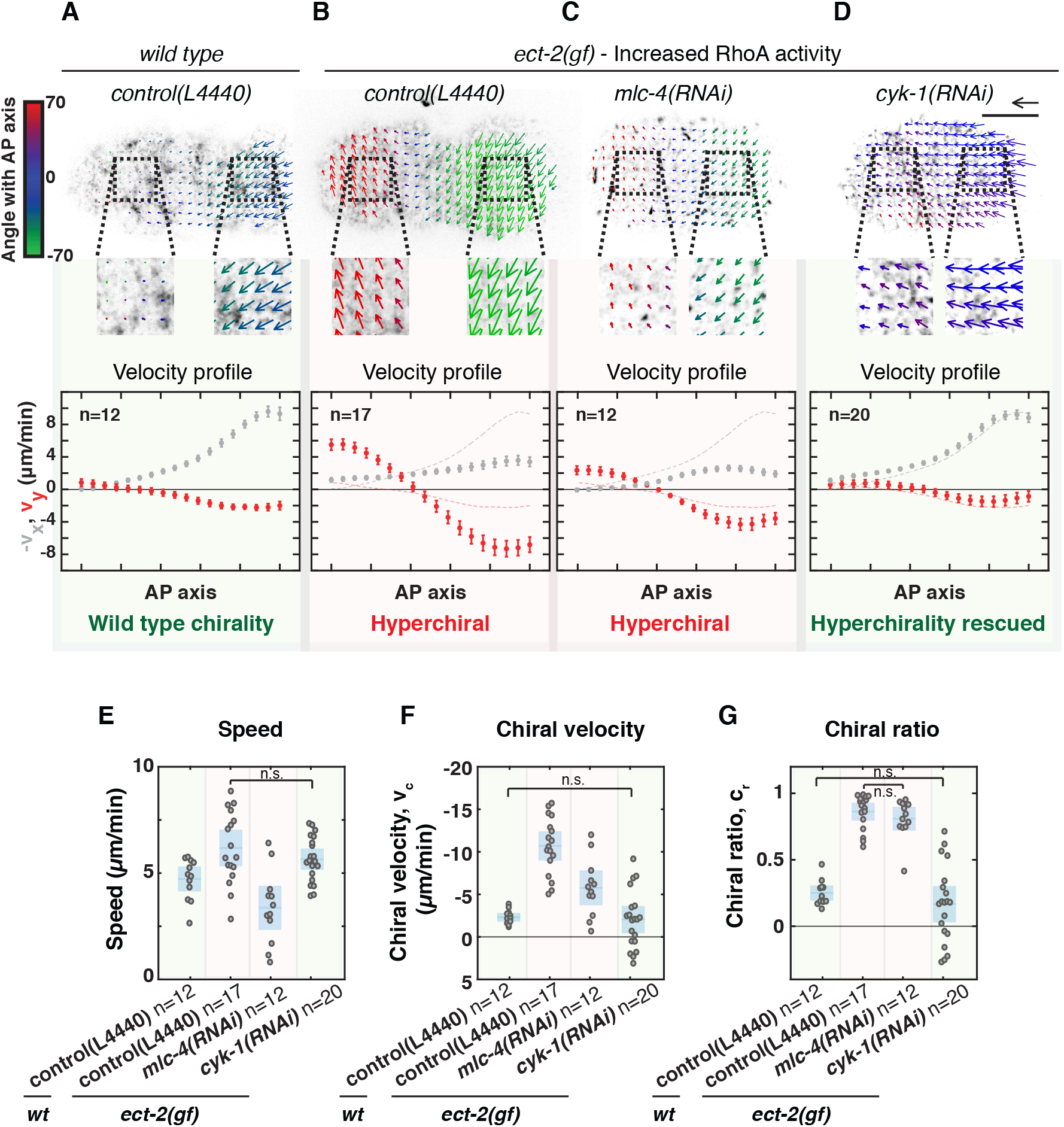
RhoA promotes chiral counter-rotating actomyosin flow via CYK-1/Formin activation. (**A**)-(**D**) Upper panels: time-averaged flow fields overlaid on a fluorescent still image of NMY-2::GFP imaged at the cortical surface of a (**A**) wild type embryo on RNAi control (L4440), an (**B**) *ect-2(gf)* mutant on RNAi control (L4440), (**C**)*ect-2(gf); mlc-4(RNAi)* and (**D**) *ect-2(gf); cyk-1(RNAi)*. Mean velocity vectors are color coded for their angle with the anteroposterior axis. Scale bar, 10 *µ*m. Velocity scale arrow, 20 *µ*m/min. Lower panels: Mean x-velocity (grey) and y-velocity (red) in 18 bins along the anteroposterior axis. Dashed lines in the plots in (**B**)-(**D**) display the mean x and y-velocity profiles in wild type. Error bars, SEM. n indicates the number of embryos. (**E**) Mean speed, (**F**)chiral velocity v_c_ and (**G**) chiral ratio c_r_ per embryo. Data points represent individual embryos. Blue stripe and area represent the mean with 95% confidence interval. Significance testing: only conditions that are not significantly different are indicated in the diagrams (n.s., p>0.05, Wilcoxon rank sum test). The flow hyperchirality (red background in all panels) of *ect-2(gf)* is rescued in *ect-2(gf); cyk-1(RNAi)* (green background in all panels), but not in *ect-2(gf); mlc-4(RNAi)* (red background in all panels).

Next, we asked how the *ect-2(gf)*-induced hyperchirality depends on the RhoA effectors Non-Muscle Myosin II and CYK-1/Formin. We first reduced Non-Muscle Myosin II activity in the *ect-2(gf)* background by performing RNAi against the Myosin regulatory light chain, *mlc-4*. Importantly, as indicated by NMY-2::GFP fluorescence levels, cortical Non-Muscle Myosin II was reduced but not fully depleted (compare Movie 14 with Movie 15). *mlc-4(RNAi)* resulted in a decrease in both the flow speed and the chiral component of the flow, when compared to *ect-2(gf)* alone (Movie 15, Figure 5C, E, F). Strikingly, the chiral ratio was indistinguishable from that observed in *ect-2(gf)* control embryos, indicating that Non-Muscle Myosin II does not influence the relative strength of chiral counter-rotation (Figure 5G). In contrast, *cyk-1(RNAi)* in *ect-2(gf)* mutants completely rescued the *ect-2(gf)*-induced hyperchirality phenotype: the chiral velocity (v_c_), the anteroposteiror velocity (v_x_) and the chiral ratio (c_r_) were all restored to wild type levels (Movie 16, Figure 5D, F, G, Supplementary figure 6A,E-F) suggesting that RhoA acts through CYK-1/Formin. Finally, we tested whether ANI-1/Anillin, another RhoA target, is required for *ect-2(gf)*-induced hyperchirality. We found that *ani-1(RNAi)* in the *ect-2(gf)* background reduced both the flow speed and the chiral ratio (Movie 17, Supplementary figure 6B-D). ANI-1/Anillin is a passive actin cross linker and therefore this result is consistent with ANI-1/Anillin playing a permissive role by controlling F-actin network integrity. Altogether, we conclude that Non-Muscle Myosin II and CYK-1/Formin play mechanistically distinct roles in the emergence of chiral flows: While myosin activity is required to drive cortical flows, CYK-1/Formin activity is required to break chiral symmetry and promote chiral counter-rotatory flows with a consistent handedness.

To determine whether RhoA signaling determines actomyosin flow chirality through CYK-1/Formin activation, or whether active RhoA and CYK-1/Formin affect flow chirality in parallel, we next performed genetic epistasis analysis. If CYK-1 and RhoA act in the same pathway, then full depletion of *cyk-1/Formin* in a RhoA hyperactive background is expected to phenocopy full depletion of *cyk-1/Formin* alone. Alternatively, if CYK-1 and RhoA control actomyoin flow chirality in parallel, a full depletion of *cyk-1/Formin* in a RhoA hyperactive background will result in an intermediate flow chirality phenotype. To discriminate between these possibilities, we next analyzed cortical flows in *cyk-1(lf)* mutants, treated with *rga-3(RNAi)*, at restrictive temperature (25°C). Consistent with our previous reports, *rga-3(RNAi)* on control embryos strongly increases actomyosin flow chirality (Movie 18, Supplementary figure 6E, [7, 36]). Conversely, *rga-3(RNAi)* on *cyk-1(lf)* mutants did not affect actomyosin flow chirality and phenocopied *cyk-1(lf)* alone (Movie 19-20, Supplementary figure 6E-F). These results demonstrate that *cyk-1/Formin* is epistatic to *rga-3*, and indicate that RhoA activity induces chiral symmetry breaking via CYK-1/Formin activation.

### CYK-1/Formin activity in active RhoA foci promotes in-plane torque generation

How are chiral counter-rotating flows generated? The active chiral fluid description is built on the assumption that the cortex generates active in-plane torques that tend to rotate the cortex clockwise (when viewed from the outside). Moreover, the cortex is organized into distinct foci, that contain active RhoA and recruit CYK-1/Formin. Therefore, we hypothesize that 1) the functional units that generate in-plane torques are active RhoA foci and 2) Formin activation in these RhoA foci determines how strong the in-plane torques will be. Given that the actomyosin cortex is a dense, highly-connected meshwork, RhoA foci are interconnected and are unlikely to be free to rotate in response to active torques. Therefore, to test these hypotheses directly, we sought to experimentally induce the formation of a single patch enriched in RhoA activity. Interestingly, in a recent study it was shown that strong compression of *C. elegans* one-cell embryos during cytokinesis resulted in a collapse of the cytokinetic ring into a cortical patch containing high levels of the RhoA target non-muscle myosin II (NMY-2) [80]. Because the cytokinetic ring forms in response to active RhoA, we hypothesized that these patches are regions of high RhoA activity. To test this, we first repeated the embryo-compression experiments and found that, in addition to NMY-2, F-actin and CYK-1/Formin are also enriched in these compression-induced cortical patches (Movie 21-23, Figure 6A-C). We conclude that indeed these patches are regions of high RhoA activity.

**Figure 6:**
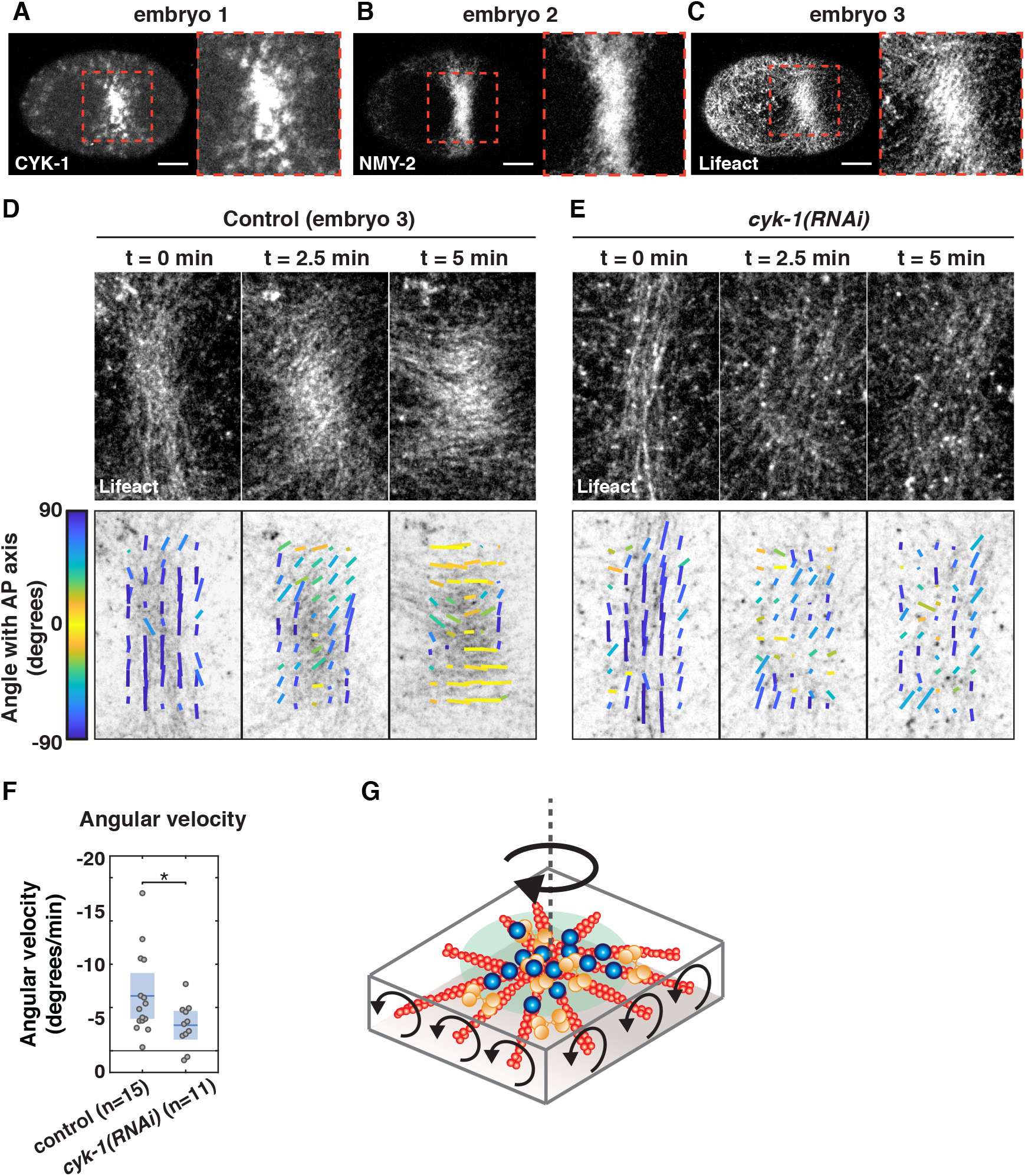
CYK-1/Formin activity in a compression-induced RhoA patch promotes clockwise reorientation of cortical F-actin. (**A**)-(**C**) Micrographs of compressed embryos that fail to divide due to a collapsed cytokinetic ring, resulting in a region enriched in (**A**) CYK-1/Formin, (**B**) NMY-2 and (**C**) F-actin. Note that the three images are derived from three different embryos. Scale bar, 10 *µ*m. (**D**-**E**) Fluorescent micrographs of cortical Lifeact-mKate2 (upper panels), overlaid with the local filament order in small templates (lower panels) of a control (**D**) and (**E**) *cyk-1(RNAi)* embryo, at the onset of patch formation (t=0 min), during patch rotation (t=2.5 min) and at the end of patch rotation (t=5 min). Both examples shown in (**D**-**E**) display an embryo with strong clockwise reorientation, with respect to the mean of the condition. Local directors are color coded for the angle with the anteroposterior axis. (**F**) Mean angular velocity in a time window of 5 minutes during reorientation of the patch in control and *cyk-1(RNAi)*. Because the timing of rotation with respect to the onset of patch formation varied among embryos, the start of the 5-minute time window was chosen such that the clockwise angular velocity was maximal (see methods). Mean with 95% confidence interval is indicated in blue. Significance testing: * represents p*<*=0.05 (Wilcoxon rank sum test). n indicates the number of embryos. (**G**) Schematic of an active RhoA patch (green area) in which F-actin (red filaments), NMY-2 (orange) and CYK-1/Formin (blue) are enriched. Due to active torque generation within the active RhoA region, the patch will reorient in a clockwise fashion (when viewed from the outside). If actin filaments are elongated by formins that are immobilized in the active RhoA patch, oppositely oriented filaments will counter-rotate, which we hypothesize to be driving the clockwise rotation of the patch as a whole (see discussion).

If regions of high RhoA activity generate in-plane torques, we expect to see clockwise reorientations of the compression-induced active RhoA patch itself and the region surrounding it. Therefore, we next analysed the average orientation of actin filaments within and around the active RhoA patch. To quantify actin filament orientation we used a previously described method that subdivides the F-actin image into small square templates, from which it extracts the mean filament orientation angle [81](Figure 6D-Elower panels). We found that at the onset of patch formation, actin filaments were mainly oriented perpendicular to the anteroposterior axis, thereby resembling a normal cytokinetic ring (Movie 23, Figure 6D, left panels). Over time cortical flow was directed towards the active RhoA patch and CYK-1, NMY-2 and F-actin accumulated in the patch (Movie 21-Movie 23). Strikingly, during growth of the patch, actin filaments slowly reoriented in a clockwise fashion with an average angular velocity of 6.34 ± 2.62°/min (Movie23, Figure 6D, F). Although the extent and the speed of filament reorientation varied substantially among embryos (Movie 23, Movie 24, Movie 25), counter-clockwise rotations were never observed (Figure 6F, Supplementary figure 7A, C, D). Therefore, these results provide direct experimental evidence for the hypothesis that active torque generating processes tend to rotate the actomyosin cortex in a clockwise fashion.

Finally, we asked whether the clockwise, in-plane rotations, are CYK-1/Formin dependent. To test this we analysed the reorientation of actin filaments within and around the compression-induced RhoA patch upon *cyk-1(RNAi)*(Movie 26, Movie 27, Figure 6E, Supplementary figure 7B). Notably, rather than performing a strong depletion, we used conditions in which ring formation and cytokinesis still occurred in uncompressed conditions, indicative of a partial *cyk-1/Formin* knock down. We found a subtle but significant reduction of actin filament reorientation upon *cyk-1(RNAi)* (2.95 ± 1.69 °/min)(Figure 6E-F, Supplementary figure 7B-D) indicating that *cyk-1/Formin* is indeed required for in-plane rotation of the active RhoA patch. Altogether, our results indicate that regions of high RhoA activity generate active torque in a CYK-1/Formin dependent manner, with consistent (clockwise) handedness.

## 5 Discussion

The work presented here shows that CYK-1/Formin, activated in cortical RhoA foci, promotes in-plane, active torque generation in the actomyosin layer and thereby facilitates left-right symmetry breaking of *C. elegans* embryos. Moreover, we find that CYK-1/Formin and Non-Muscle Myosin 2 (NMY-2) play mechanistically distinct roles: While NMY-2 is necessary to drive the emergence of cortical actomyosin flows, CYK-1/Formin drives chiral symmetry breaking of the flow. Altogether, our work provides mechanistic insight into the role of Myosins and Formins in organismal left-right symmetry breaking.

How could CYK-1/Formin activity in RhoA foci promote in-plane torque generation with clockwise handedness? One attractive possibility is that the formins themselves generate the active torque while elongating actin filaments. Due to the helical nature of filamentous actin, rotationally constrained Formins rotate the actin filaments that they are elongating *in vitro* [44]. If CYK-1/Formins in active RhoA foci are indeed immobilized and rotationally constrained, the actin filaments that they are elongating in opposite directions will counter-rotate (Figure 6G). In turn, this would result in a large scale torque with a clockwise handedness if a difference in friction between two sides of the cortical layer is present, e.g. the cortex experiencing lower friction with the membrane than with the cytosol. The rotating active RhoA patch could be compared to a Segway (or a military tank, but the authors prefer pacifist allegories) rotating on the spot. In this analogy, the wheels represent oppositely oriented actin filaments and their counter-rotation will result in an in-plane rotation of the structure as a whole (RhoA patch or Segway).

Following this proposition, we expect active RhoA foci to be at the center of polar actin asters (Figure 6G). To test this we have performed SIM-TIRF super-resolution microscopy on embryos producing Lifeact-mKate2 and NMY-2/Myosin::GFP. Although actin filaments often appear isotropically oriented, we clearly observe aster-like topology around many of the active RhoA foci (Movie 28-29, Supplementary figure 8). Notably, it has been shown that *in vitro* reconstituted contractile actin networks have the propensity to self-organize in polar asters, with Myosins and actin plus-ends at the aster center [82]. Even though the aster-like structures in the *C. elegans* cortex are far from ideal asters they could, on average, still facilitate in-plane torque generation. Moreover, for a net torque with consistent handedness an aster-like topology might not be necessary. Any organization in which densely clustered Formins, that are elongating actin filaments in opposite orientations, could be sufficient to generate in-plane rotations. Consistent with this proposition are earlier reports in cultured cells showing that Formins anchored at peripheral adhesion sites, that are elongating actin filaments towards the cell center, are driving chiral in-plane swirling [25, 27].

Although the Formin clustering proposition provides a possible mechanism for chiral symmetry breaking, several open questions remain. First, how could actin filaments in a highly cross-linked actomyosin cortex rotate? Previous *in vitro* studies have shown that, when both the Formin and the actin filament are rotationally constrained, Formins stall actin polymerization [83, 84] and this would attenuate their ability to generate torques *in vivo*. Second, what is the role of myosins in active torque generation? Because NMY-2 activity is required for actomyosin flows to emerge, Formin activity itself is not sufficient to generate chiral counter-rotatory flows. NMY-2 may therefore facilitate Formin-driven active torque generation. Interestingly, in a recent study Noselli and colleagues reported that the Formin DAAM is required for Myosin 1D (Myo1D)-driven left-right patterning of the *Drosophila* body plan [47]. Their biochemical analysis revealed that Myo1D and DAAM physically interact, suggesting that Myosins and Formins could influence their activities reciprocally. Although NMY-2 is a different type of Myosin, our finding that NMY-2 and CYK-1/Formin reside in the same active RhoA foci, could point out common regulatory mechanisms. Notably, numerous Formins have been shown to be mechanosensitive and the Diaphanous-like Formin mDia1 speeds up polymerization when pulling force is exerted on the actin filament [84, 85, 86]. In the *in vivo* cortex, NMY-2 motors generate contractility and hence are likely candidates for providing pulling forces. The role of NMY-2 might therefore be to reactivate actin polymerization by stalled Formins, thereby recovering the potential to generate active torques. Future studies may be aimed at understanding how the complex, and highly regulated, molecular interplay between Formins and Myosins facilitates active torque generation.

The work presented here, together with our previous work [7, 36], shows that Formin dependent chiral counter-rotating actomyosin flows drive cell division skews that are essential for left-right patterning in *C. elegans* embryos. Chiral counter-rotations of the same handedness have been observed in Xenopus embryos [33], and both Xenopus and snail left-right patterning depends on Formins [14, 15, 46]. However, it remains unclear whether chiral counter-rotating actomyosin flows drive chiral morphogenesis in other animal species. Interestingly, it has been shown that the snail Diaphanous-like formin, LsDia, is the critical left-right determinant that sets the dextral handedness of the body plan [15]. As in *C. elegans* embryos, left-right symmetry breaking in snails occurs through a cell division skew in the 4-cell embryo and it is therefore an attractive hypothesis that Formin-dependent chiral counter-rotating flows are also involved in chiral patterning of snails. Altogether, Formin-dependent chiral counter-rotating flows may drive cell division skews throughout the animal kingdom.

## 6 Materials and Methods

### Worm strains and culture

*C. elegans* strains were cultured using standard culture conditions [87]. The N2 Bristol strain was used as non-transgenic wild type control. Strains were maintained at 20°C, apart from strains carrying *cyk-1(or596ts)*, which were maintained at 15°C. Alleles and transgenes used in this study are: LGI, *nmy-2(cp8[nmy-2::GFP])*[55], *nmy-2(cp52[nmy-2::mkate2])*[88]. LGII, *gesSi48[mCherry-ani-1(AHPH)]* (this study), *xsSi5[GFP-ani-1(AHPH)]* [71], *ttTi5605* [89], *ect-2(zh8)*[79]. LGIII, *cyk-1(or596ts)*[54], *cyk-1(ges1[cyk-1::GFP])*[81], *unc-119(ed3)*[90]. LG unknown, *gesIs002[Lifeact::tagRFP]* [70], *gesIs003[lifeact-mKate2]* [81], *weIs21[mCherry::tubulin]* (Ahringer lab).

### Molecular cloning and generation of transgenes

In order to generate the constitutively active *cyk-1/Formin* construct (CA-CYK-1/Formin) we first codon optimized the *cyk-1/Formin* cDNA, covering residues 700-1437, using the ‘*C. elegans* codon adapter’ online tool [91], made the construct resistant to RNAi of the original gene, avoided splice sites in the codon region and introduced one artificial intron. The resulting *cyk-1/Formin* cDNA was synthesized (Genscript). In addition, the *mex-5* promoter and the *tbb-2* 3’UTR were obtained from pJA245 and pCM1.36 respectively [92], a Pleckstrin Homology domain was obtained from *Pwrt2::PH-GFPcoLov* [93] and a *GFP*, containing *smu-1* introns to prevent germline silencing, was obtained from pCFJ1415 (Addgene) [94]. The final construct, *Pmex-5::PH-GFP-cyk-1(700-1437)::tbb-2 3’UTR* in the pCFJ150 backbone [89] was made using HiFi cloning (New England Biolabs). Notably, we included a 12 residue linker (SGM followed by 3x GGS) at the C-terminal end of GFP. Moreover, due to intermediate subcloning steps, an attB1 site (resulting in TSLYKKAGS in the fusion protein) was located in between the linker and the start of *cyk-1(700-1437)*. As a negative control we replaced *cyk-1(700-1437)* by a LOV2 domain derived from *Pwrt2::PH-GFPcoLov* [93], which should not have any effect on actomyosin activity. For transient expression, the resulting plasmids were diluted to 20 ng/*µ*l in deionized water and injected into the gonad of young adults that expressed *lifeact-mKate2*. The injected worms were grown at 25°for 6-7 hrs and thereafter their progeny was imaged at room temperature (22°C).

In order to generate the *mCherry::ani-1(AH-PH)* transgene, the portion of *ani-1* as used in [71], containing the AH-PH domains, was PCR amplified from MG617 [71] worm lysate and cloned into the pDONR221 vector using BP clonase (Gateway technology, Thermofisher). *Pmex-5::mCherry::ani-1(AH-PH)::tbb-2* 3’UTR in pCFJ150 was obtained by mixing *ani-1(AH-PH)* in pDONR221, pJA281 (containing *Pmex-5::mCherry*), pCM1.36 (containing *tbb-2* 3’UTR), and pCFJ150 (containing *Cbr-unc-119* rescue module and homology arms for Mos-mediated single copy integration into the *ttTi5606* site)[89, 92] with LR clonase II+ (Gateway technology) followed by transformation into competent cells. Transgenesis was done using Mos single copy insertion into the *ttTi5605* site as previously described [89]. After verification of the insertion, the strain was outcrossed with N2 wild type 2 times and subsequently crossed with *cyk-1(ges1[cyk-1::GFP])*[81].

### RNA interference

All RNAi treatments were performed by feeding as previously described [95]. Briefly, NGM agar plates containing 1 mM isopropyl-*β*-D-thiogalactoside and 50 *µ*g ml-1 ampicillin were seeded with bacteria expressing dsRNA targeting the gene of interest. The L4440 control clone, *rga-3, ani-1/Anillin, cyk-1/Formin* RNAi clones were obtained from the Ahringer RNAi library (Source BioScience) [96]. The *mlc-4* and *ect-2* RNAi clones were obtained from the Hyman lab. For the titration of the *cyk-1(RNAi)* (Figure 2), L4 larvae or young adults were grown on feeding RNAi plates at 20°C for different durations ranging from 6 to 24 hrs. For *cyk-1(RNAi)* treatments shorter than 9 hrs, L4 worms were first grown for 20-24 hrs on OP50, and then transferred to feeding RNAi plates where they were grown for an additional 6-7 hrs or 7.5-8.5 hrs. *cyk-1(RNAi)* treatments in the range of 9-24 hrs were done by transferring L4 animals to *cyk-1/Formin* feeding RNAi plates for 9-10 hrs, 10-11 hrs, 12-13 hrs, 15-16 hrs, 17-18 hrs, or 22-24 hrs. *rga-3(RNAi)* was performed by transferring L4 larvae to feeding RNAi plates and growing them for 49-51 hrs at 20°C. For the CYK-1/Formin::GFP speckle microscopy experiments 24-26 hr *rga-3(RNAi)* treatments were performed. *ani-1(RNAi)* on the *ect-2(zh8)* mutant was done by transferring L2-L3 larvae to feeding RNAi plates for 48-50 hrs at 20°C. *cyk-1(RNAi)* on the *ect-2(zh8)* mutant was done by transferring L2-L3 larvae to feeding RNAi plates for 40-41 hrs (mild) or 52-54 hrs (strong), keeping them at 20°C. *mlc-4(RNAi)* on the *ect-2(zh8)* mutant was done by transferring L4 larvae to *mlc-4* feeding RNAi plates for 24-25 hrs at 20°C. We note that the duration of the RNAi treatment of *ect-2(zh8)* mutants is increased when compared to control worms, which may be because of a reduced sensitivity to RNAi or higher baseline levels of cortical NMY-2 and CYK-1/Formin activity in this mutant. As a negative control for the RNAi treatments in the experiments assaying cortical flows and cortical protein levels, worms were fed bacteria containing empty vector (L4440) for the same number of hrs. For the analysis of the collapsed cytokinetic ring upon *cyk-1(RNAi)* treatment, L4 worms were transferred to feeding RNAi plates and grown at 25°C for 24 hrs.

### Image acquisition

Apart from the CYK-1/Fromin::GFP speckle microscopy, all imaging was done using spinning disk confocal microscopy on a Zeiss Axio Observer Z1 equipped with a Yokogawa CSU-X1 scan head, a 488 nm and 561 excitation laser, a C-Apochromat 63X/1.2 NA W objective, a Hamamatsu ORCA-flash 4.0 camera (final pixel size 0.106 *µ*m), operated by Micromanager software. Imaging of *cyk-1/Formin* mutants and the accompanying controls was performed at 25°C using a CherryTemp temperature-control stage (Cherry Biotech). Imaging of the control *nmy-2::GFP* strain at 16°C, 22 °C and 25 °C was also performed using the CherryTemp temperature-control stage. The rest of the imaging in this study was performed at room temperature (22 °C).

CYK-1/Formin::GFP speckle microscopy videos were recorded using spinning disc comfocal microscopy on a Nikon Ti-E inverted microscope equipped with a Yokogawa CSU-X1 scan head, a 488 nm excitation laser, a CFI Apochromat 100X/1.49 NA Oil TIRF objective, an Andor iXon Ultra camera, a Nikon perfect focus hardware autofocus system (final pixel size 0.068 *µ*m), operated by Nikon Elements software.

2D SIM-TIRF imaging of embryos expressing Lifeact-mKate2 and NMY-2::GFP, was performed on a GE-Deltavision OMX equiped with a 60X/1.42 NA Oil objective and a pco edge 5.5 sCmos camera operated by OMX Softworks software. Excitation was done using 488 and 568 nm, 3 angles of illumunation with 3 phases each, linespacing 488: 0.182 *µ*um, linespacing 568: 0.217 *µ*um. Reconstruction was done in OMX Softworks software with a base background subtraction of 70, and a baseWienerSNR of 0.001. After image reconstruction bleach correction was perfomed in Fiji.

For imaging early polarizing flows, and cortical protein localization during polarizing flows, worm embryos were dissected and mounted in M9 on 2% agarose pads. Subsequently, at each timepoint (unless stated differently) a small z-stack of 2 slices with 0.5 *µ*m spacing was made covering the cortical surface. A maximum intensity projection of this stack at each timepoint was made prior to analysis. For cortical flow measurements during polarization, side-polarized embryos (in which the paternal pronucleus first appeared at a position distant from the embryo pole) were excluded from the analysis. Embryos were imaged starting from the beginning of polarizing flows until pronuclear meeting. To image cortical flows during anteroposterior polarization of embryos producing NMY-2::GFP 5 second intervals were used. For imaging CYK-1/Formin::GFP during polarizing flow in control and upon *rga-3(RNAi)* embryos were imaged with 10 second intervals. For dual color imaging of NMY-2::mKate2 and CYK-1/Formin::GFP, 5 second time intervals were used for NMY-2::mKate and, in order to prevent photobleaching, 20 second intervals were used for CYK-1/Formin::GFP. For dual color imaging of Lifeact-mKate2 and CA-CYK-1/Formin or control (PH-GFP-LOV2) during anteroposterior polarization 5 second intervals were used for Lifeact-mKate2 and, in order to prevent photobleaching 30 second intervals for CA-CYK-1/Formin and PH-GFP-LOV2. For dual color imaging of CYK-1/Formin::GFP and mCherry::ANI-1(AH-PH) during polarizing flows 5 second intervals were used for both channels.

Imaging of mCherry-tubulin during the cell division skew of ABa and ABp in control and *cyk-1/Formin* mutant embryos was performed as previously described [36]. Briefly, z-stacks composed of 21 slices 1 *µ*m apart, covering the mitotic spindles of ABa and ABp, were captured every 5 seconds starting from metaphase until the end of cytokinesis.

For CYK-1/Formin::GFP speckle microscopy, worm embryos were dissected and mounted in M9 on 2% agarose pads. The Nikon perfect focus system was used to keep the focal plane constant throughout imaging. Videos of the cortical surface were made with 50 ms time intervals.

To image one-cell embryos during compression-induced collapse of the cytokinetic ring, embryos were dissected on a cover glass in a drop of 65% Shelton’s Growth Medium (SGM) with 35% Fetal Calf Serum containing 10 *µ*m polystyrene spacer beads (Polysciences). Subsequently a glass slide was lowered over it such that the embryos were confined between the cover glass and the glass slide. The sample was sealed using valap to prevent evaporation of the buffer during imaging. Every 3 seconds, 3 z-slices with 0.4 *µ*m spacing covering the actomyosin cortex were captured of NMY-2::GFP and Lifeact::tagRFP or CYK-1/Formin::GFP. A maximum intensity projection was made of the z-stacks covering the actomyosin cortex prior to further analysis.

### Image analysis

Cortical actomyosin flows were quantified using PIV, similar to previously described [7]. Briefly, 3-step PIV (with a final box size of 18×18 pixels on a grid with 9 pixel spacing) was done using an open source Matlab package (PIVlab) [97]. The obtained flow fields were rotated to align the anteroposterior axis horizontally (anterior left, posterior right in all cases), averaged over time (100 seconds during polarizing flows), and finally, spatial averages in a stripe of 120 pixels high (12.72 *µ*m) subdivided into 18 bins along the AP axis were calculated. In order to obtain the v_c_, flow speed and chiral ratio, velocity vectors in anterior bins (number 3-6) and in posterior bins (number 13-16) were first averaged to obtain a mean anterior flow velocity vector and a mean posterior flow velocity vector. In order to obtain the chiral counter-rotating velocity v_c_ the y component of the anterior velocity vector was subtracted from the y component of the posterior velocity vector. To obtain the flow speed the mean of the anterior and posterior flow magnitude was computed. The chiral ratio, c_r_ is defined as 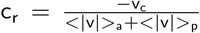. To obtain the flow decay length we first extracted the mean flow field in the central part (e.g. the central 55%) of the horizontal stripe along the anteroposterior axis. Subsequently, the autocorrelation of the x component of the velocity vectors along the anteroposterior axis was computed. The decay length was defined as the length along the anteroposterior axis at which the normalized autocorrelation dropped below 0.5.

In order to quantify the typical cortical fluorescence intensity of CA-CYK-1/Formin, we measured cortical GFP levels in a masked region covering the embryo. However, diffuse cortical GFP signal was indistinguishable from background fluorescence, and CA-CYK-1/Formin signal was only observed in sparsely localized speckles at later stages of early flow. Therefore, we quantified the summed fluorescence of GFP in speckles whose intensity exceeded that of the mean fluorescence plus 3 standard deviations at the end of the early flow phase. For the cortical flow analysis in Fig. 1 we only included embryos that had a summed fluorescence within the specified range.

To obtain mean cortical concentration profiles of NMY-2::GFP, NMY-2::mKate2 and CYK-1/Formin::GFP, the same time points were used as for computing the mean flow profile. Movies were first rotated (applying bilinear pixel interpolation) to align the anteroposterior axis horizontally (anterior left, posterior right). In each time point, the mean pixel value in 18 spacial bins along a anteroposterior stripe (same as for obtaining flow profiles) was computed and, subsequently, a time-averaged anteroposterior concentration profile was obtained for each embryo. Background subtraction for NMY-2::GFP and NMY-2::mKate2 was performed for each embryo by subtracting the mean pixel value of a region outside of the embryo. From the anteroposterior profiles, we obtained the typical cortical levels for each embryo by taking the mean fluorescence level of bins 9-11. For the quantification of cortical CYK-1/Formin::GFP we found that the GFP signal was close to that of autofluorescence. Therefore, in addition to the background level fluorescence outside of the embryo, we subtracted the mean autofluorescence signal obtained by imaging non-transgenic N2 zygotes using the same imaging conditions. Cytoplasmic signal of Lifeact-mKate2 and of CYK-1::GFP was measured in a midplane slice 10 *µ*m below the cortical surface.

To quantify the co-localization of mCherry::ANI-1(AH-PH) and CYK-1/Formin::GFP we used the first 75 seconds (15 time points) of the polarizing flow phase. Images were binned 3×3 (to get to a binned pixel size of 318 nm), and for each time point a mask was made covering the embryo. A pixel-by-pixel Pearson correlation coefficient was computed using all the binned pixels within the mask in each time frame. As a negative control for correlations, in each time frame, the pixel positions in one of the binned channels were scrambled randomly 100 times. For both CYK-1/Formin::GFP and mCherry::ANI-1(AHPH) the fluorescence signal was not far above autofluorescence. Therefore, as an additional control, non-transgenic wild type (N2) embryos were imaged using the same conditions, and the same analysis was performed. Although autofluorescence weakly correlated, this was substantially less than the correlation found for CYK-1/Formin::GFP and mCherry-ANI-1(AHPH).

For analysis of the CYK-1/Formin::GFP speckles, programming, data extraction, wrangling, calculations and plotting were done using Python 3.7 with standard scientific libraries (Oliphant, 2007; Jones et al.; Hunter, 2007; Millman and Aivazis, 2011). Embryo masks were generated by segmenting the first unbleached frame, using a mean threshold followed by manual inspection and correction when necessary. The first 1000 frames of each movie were discarded. Candidate 2D fluorescent peaks were detected and tracked using Trackpy [98], and peaks outside embryo masks were discarded. Peaks in consecutive frames, and for up to 5 discontinuous frames, were considered the same if they re-appeared in a neighborhood with radius of 3 pixels. A Gaussian-Process Classifier (GPC) trained with a sample set of manually classified images was used to distinguish formin foci from spurious peaks. Only peaks where at least one frame-appearance had a GPC probability of at least 0.5 were kept. This approach yielded results qualitatively consistent with discarding peaks that were detected in a single frame only.

The analysis of the ABa and ABp spindle skews during cytokinesis were performed as previously described [36]. Briefly, the movies were first rotated using the clear volume plugin [99] in FIJI and projected such that the spindle skew occurred in the projection plane. The cell divsion skew angle was subsequently defined as the difference between the initial orientation of the spindle and the final orientation upon completion of cytokinesis.

For quantifying the typical angle of actin filaments within the compression-induced region of high RhoA activity, we first rotated the Lifeact-tagRFP movies to align the anteroposterior axis horizontally anterior left, posterior right). Subsequently, we manually defined a rectangular region of interest of 100 pixels by 200 pixels (10.6 *µ*m by 21.2 *µ*m) covering the region of high RhoA activity. This region of interest was subdivided into small square templates of 40×40 pixels, using a grid with 20 pixel spacing. In each 40×40 pixel template, the actin filament orientation was computed using a method that we previously developed [81]. Importantly, this indirect method does not segment individual filaments (which is challenging due to high filament density) but is based on Fourier Transform analysis of the fluorescent image intensities and quantifies a mean filament orientation for each template. Subsequently, we computed the mean filament angle over all the templates in each time point. Importantly, because we have no information on the polar orientation of filaments the orientation angles have a periodicity of *π* instead of 2*π*. Therefore, the measured angles were first multiplied by 2, followed by calculation of the mean angle using the circular statistics toolbox for matlab [100], and then divided by 2 to get the final mean angle. The mean angle was calculated in each time frame starting from the onset of patch formation (manually defined) until 9 minutes thereafter. Because the raw traces of angles were noisy, we computed a moving average with a one-minute sliding window. In order to obtain the angular velocity (plotted in Fig. 6E) we extracted the difference in mean orientation angle over a time window of 5 minutes. Because the timing of rotation with resp[ect to onset of patch formation varied substantially among embryos, we chose the start of the 5 minute time window such that the clockwise angular velocity was maximal. Therefore, the time window with respect to the onset of patch formation varied among embryos.

### Fitting the hydrodynamic model to experimental measurements

To obtain measures of active tension and active torque generation, hydrodynamic equations of motion were fit to the experimentally measured vx and vy profiles along the AP axis as previously described [7]. We used the measured anteroposterior myosin distribution as a proxy for the distribution of active tension and active torque. In order to obtain error estimates for the fit parameters (the tension and torque coefficients as well as the hydrodynamic length), for the cortical CYK-1/Formin concentration and for the fitted velocity profiles, we performed bootstrapping, using 100 bootstrap samples with replacement for each of the conditions.

## 7 Other details

## Acknowledgements

We thank Julie Canman for sharing the *cyk-1(or596ts)* mutant, Bob Goldstein for sharing *nmy-2(cp8[nmy-2::GFP])* and *nmy-2(nmy-2(cp52[nmy-2::mKate2)*, Tony Hyman for sharing the *mCherry-tubulin* strain, *mlc-4* and *ect-2* RNAi clones, Martin Harterink and Sander van den Heuvel for sharing the PH-GFP-LOV2 plasmid, the CGC for providing the *ect-2(gf)* mutant and *xsSi5[GFP-ani-1(AH-PH)]*, Addgene for the pJA281, pJA245, pCM1.36, pCFJ150, pCFJ1415 plasmids, Julie Ahringer and Source BioScience for providing the L4440, *rga-3* and *cyk-1/Formin* RNAi clones. We also thank Friederike Thonwart for assistance with molecular biology, GE-Deltavision and its representatives for having the Deltavision OMX SIM-TIRF system available for the Woods Hole physiology course 2018 and thank Sylvia Hurlimann for capturing the SIM-TIRF movies during this course. Furthermore, we thank Jonas Neipel for valuable discussion on the hydrodynamic theory. We thank Anne Grapin-Botton, Arghyadip Mukherjee, Ján Sabó, Jonas Neipel and Zdeněk Lánsk’y for critical reading of the manuscript.

## Author Contributions

T.C.M. and S.W.G. conceptualized the work, analyzed the data and wrote the manuscript together. T.C.M. performed and analyzed the experiments and applied chiral active fluid theory. J.B.G. performed and analyzed the compression experiments with help from T.C.M and P.G. P.Q.C. performed the co-localization analysis, performed the speckle microscopy experiments, and extracted and analyzed the speckle microscopy data with help of P.S.. L.P. performed and analyzed the cell division skew experiments with help from T.C.M. S.Y. helped with the hyperchirality rescue experiments.

## Correspondence

Correspondence and requests for materials should be addressed to S.W.G.

## 10 Supplementary notes

### The chiral velocity is dependent on temperature and genetic background

We found that the chiral velocity in control embryos displayed quantitative differences with our previously reported value [7]. This may have several causes. First, in our previous study we used a strain expressing additional copies of fluorescently labeled *nmy-2*, while the strain used in this study expresses endogenously labeled *nmy-2*. In general, we note that transgenic strains with different backgrounds can display quantitative differences in chiral cortical flow dynamics. Second, in our previous study, all imaging was performed at room temperature, while the controls for the *cyk-1/Formin* mutant were imaged at 25°C. To systematically determine the effect of temperature on actomyosin flow chirality, we imaged cortical flows in control embryos at 16°C, 22°C and 25°C side by side. While the flow speed was only mildly affected, both the chiral velocity and the chiral ratio increased with increasing temperature (Supplementary figure 2) indicating that actomyosin flow chirality is temperature dependent. We conclude that both genetic background and temperature have subtle effects on actomyosin flow chirality.

### High levels of CA-CYK-1/Formin lead to complex cortical flow behaviour

Upon introduction of high levels of CA-CYK-1/Formin the cortex displayed complex behavior in which both flow direction and actin filament orientation reoriented multiple times during the polarizing flow (Movie 4). Moreover, the chiral ratio in these embryos was increased with respect to control injections in 4 embryos, unchanged in 3 embryos and it was reversed in 2 embryos (Supplementary figure 1E). We hypothesize that an excess of actin filaments in the active cortex may lead may turn the actomyosin cortex into an active nematic fluid, which has been observed in microtubule-based *in vitro* systems. In such a class of active fluids, internal activity of the fluid drives the emergence and disappearance of topological defects, which in turn, leads to complex flow orientations [101, 102]. The repeated reorientations of flows and actin filaments, accompanied by the reversal of the chiral ratio (in 2 out of 9 CA-CYK-1_high_ embryos) may be explained by the cortex behaving as an active nematic fluid.

## 12 Supplementary Figures

**Supplementary figure 1:**
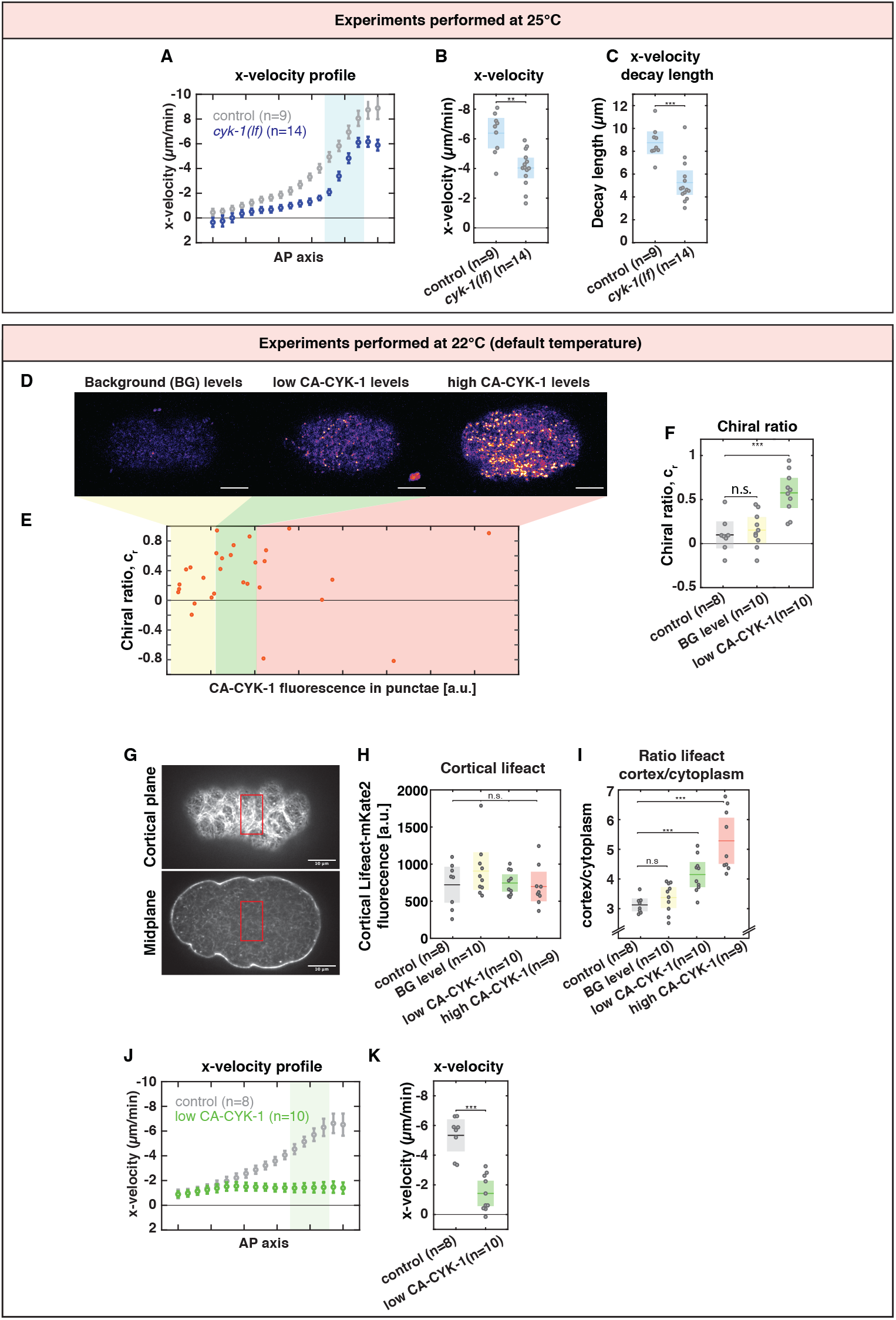
(**A**) Mean anteroposterior velocity (x-velocity) in 18 bins along the anteroposterior axis averaged over embryos for control (grey) and *cyk-1(lf)* (blue). Light blue area marks bin number 13-16 corresponding to the posterior ROI used to compute the mean x-velocity. (**B**) Spatially averaged mean x-velocity in a posterior ROI, *<* v_x_ >_p_, as indicated in (**A**). (**C**) Decay length of the x-velocity along the anteroposterior axis. The decay length was obtained by computing the normalized cross correlation of flow velocity vectors along the anteroposterior axis. We defined the decay length as as the length in */*mum at which the normalized cross correlation reached 0.5. (**D**) Cortical GFP fluorescence at the end of polarizing flow in embryos derived from *ca-cyk-1* injected parents. Lookup table: fire. Scale bar: 10 */*mum. (**E**) Chiral ratio c_r_ plotted over punctate CA-CYK-1/Formin levels. Data points represent individual embryos. Because cortical fluorescence at the onset of polarizing flow did not exceed background levels, CA-CYK-1/Formin levels were quantified at the end of polarizing flow, when CA-CYK-1/Formin appeared in distinct cortical punctae (**D**, middle and right panel). Punctate CA-CYK-1/Formin levels were measured by taking the sum of the pixel intensities that exceeded the mean cortical GFP fluorescence by three standard deviations. Because high levels of CA-CYK-1/Formin perturbed the cortex dramatically, only embryos producing CA-CYK-1/Formin within a given range (orange rectangle) were included in the analysis in (**G**)-**H** and main figure 1). (**F**) Chiral ratio c_r_ in individual embryos upon control injection, embryos derived from *ca-cyk-1* injection producing undetectable CA-CYK-1 levels (BG level) and embryos derived from *ca-cyk-1* injection producing low levels of CA-CYK-1 as indicated in (**E**). In the diagram, the data for control injection and the low CA-CYK-1 are the same as in main figure 1L. (**G**) Micrographs of the cortical plane (top) and midplane (bottom) of a control embryo producing Lifeact-mKate2. Red rectangles indicate the regions used for the measurements shown in (**H**)-(**I**). (**H**) Cortical levels and (**I**) cortex-to-cytoplasm ratio of lifeact-mKate2 in control and upon introduction of different levels of CA-CYK-1/Formin (color code as in E). Due to embryo-to-embryo variability in the Lifeact-mKate2 levels, no difference in absolute concentration of cortical Lifeact-mKate2 was observed. (**J**) Mean anteroposterior velocity (x-velocity) in 18 bins along the anteroposterior axis averaged over embryos derived from parents injected with control (grey) or *ca-cyk-1* plasmid (blue). **K** Spatially averaged mean x-velocity in a posterior ROI, *<* v_x_ >_p_, as indicated in (**J**, light blue rectangle, bin number 13-16). Experiments described in (**A**)-(**C**) were performed at 25°C. Experiments described in (**D** - **K**) were performed at 22°C. Significance testing: n.s represents p> 0.05, ** represents p*<*=0.01, *** represents p*<*=0.001 (Wilcoxon rank sum test). n indicates the number of embryos.

**Supplementary figure 2:**
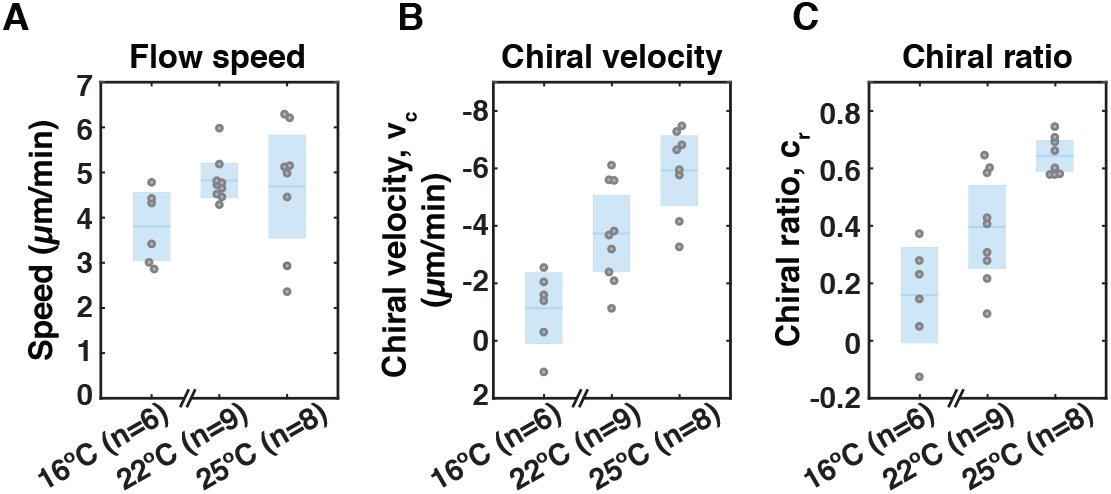
(**A**) Cortical flow speed, (**B**) chiral velocity and (**C**) chiral ratio of control worms expressing endogenously labeled *nmy-2::GFP* at 16 °C, 22 °C and 25 °C. Data points represent individual embryos. Blue boxes represent mean with 95 % confidence interval.

**Supplementary figure 3:**
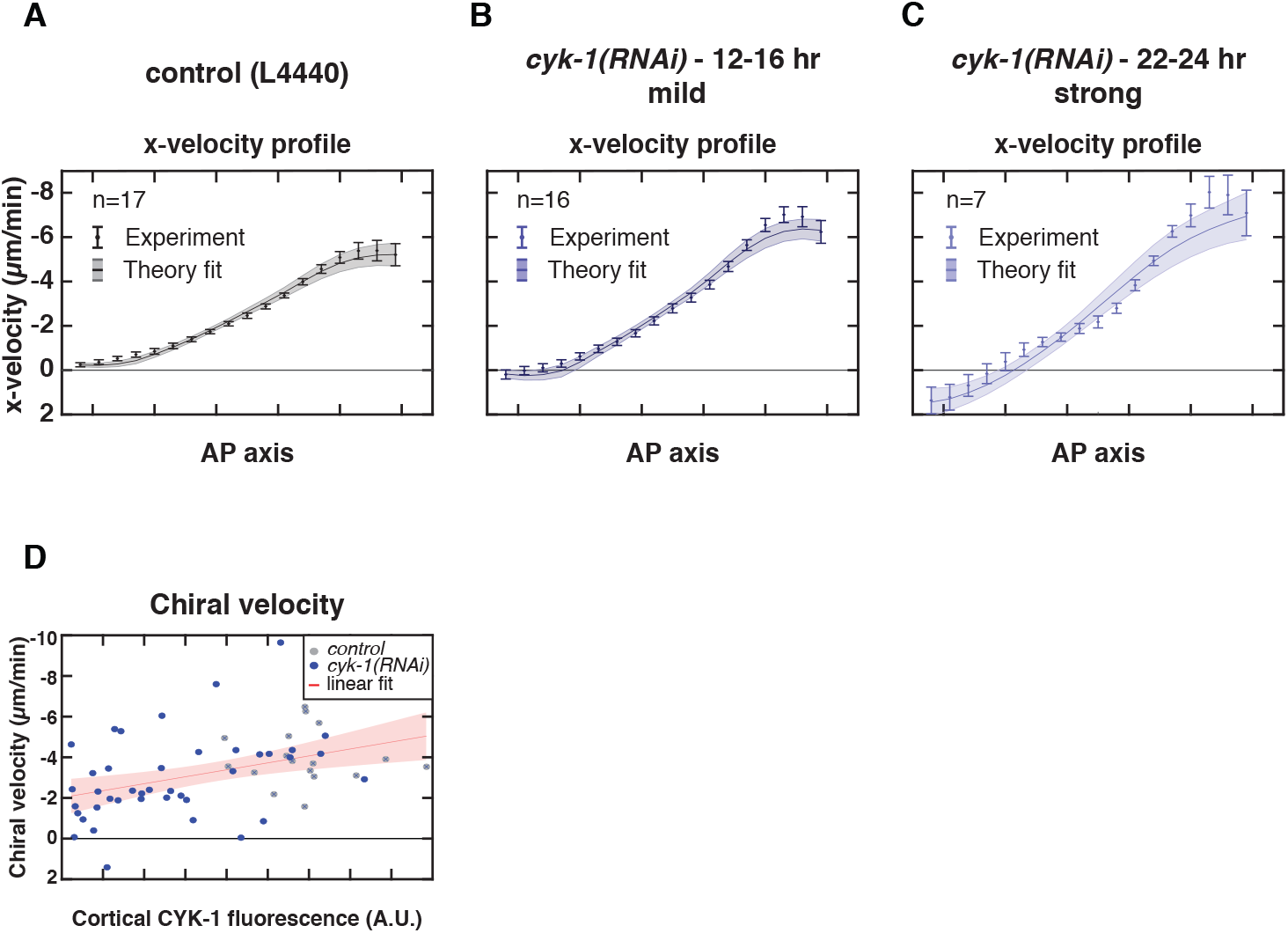
Mean x-velocity profile in 18 bins along the anteroposterior axis for (**A**) control (L4440),(**B**) mild *cyk-1(RNAi)* (12-16 hrs) and (**C**) strong *cyk-1(RNAi)* (22-24 hrs). Circles with errorbars are the experimentally measured y-velocities averaged over embryos, with SEM. Solid line shows the mean x-velocities derived from fitting the hydrodynamic model to 100 bootstrap samples with replacement. The shaded region displays the standard deviation of the mean, derived from the bootstrapping. n indicates the number of embryos. (**D**) Chiral velocity, v_c_, plotted over the measured cortical CYK-1/Formin::GFP fluorescence in control (L4440, grey) and upon increasing strength of *cyk-1(RNAi)* (blue). Data points represent individual embryos. Red line with shaded region shows a linear fit with 95% confidence bounds. The chiral velocity correlates significantly with cortical CYK-1/Formin::GFP (Spearman’s *ρ*=-0.44, p*<*0.0002).

**Supplementary figure 4:**
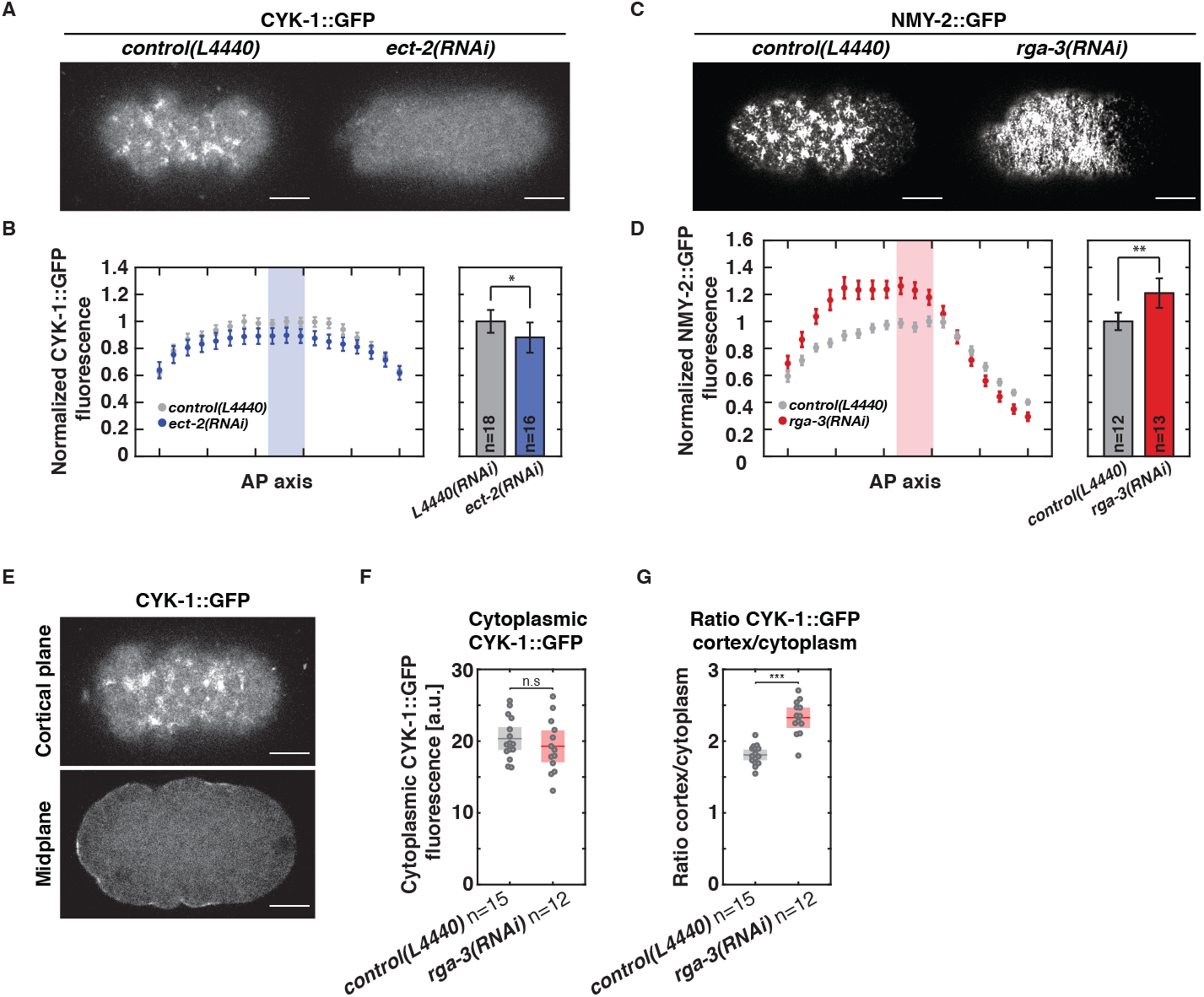
(**A**) Fluorescent micrographs of the cortical plane of embryos producing CYK-1/Fromin::GFP captured during polarizing flows in control (L4440, left) and upon *ect-2(RNAi)* (right). (**B**) Left panel: Profile of the mean cortical CYK-1/Formin::GFP fluorescence in 18 bins along the anteroposterior axis in control (L4440, grey) and upon *rga-3(RNAi)*(blue). Fluorescence levels were normalized to the mean levels in bins 9-11 (light blue rectangle). Error bars, SEM. Right panel: bar diagrams displaying the mean cortical CYK-1/Formin::GFP fluorescence measured in bins 9-11 (light red rectangle in the left panels). Error bars, 95% confidence interval. (**C**) Fluorescent micrographs of the cortical plane of embryos producing NMY-2::GFP captured during polarizing flows in control (L4440, left) and upon *rga-3(RNAi)* (right). (**D**) Left panel: Profile of the mean cortical NMY-2::GFP fluorescence in 18 bins along the anteroposterior axis in control (L4440, grey) and upon *rga-3(RNAi)*(red). Fluorescence levels were normalized to the mean levels in bins 9-11 (light red rectangle). Error bars, SEM. Right panel: bar diagrams displaying the mean cortical NMY-2::GFP fluorescence measured in bins 9-11 (light red rectangle in the left panels). Error bars, 95% confidence interval. (**E**) Micrographs of the cortical plane (top) and midplane (bottom) of a control embryo producing CYK-1/Formin::GFP. (**F**) Cortical levels and (**G**) cortex-to-cytoplasm ratio of CYK-1/Formin::GFP in control and upon *rga-3(RNAi)*. Significance testing: * represents p *<*=0.05, ** represents p *<* =0.01, *** represents p *<* =0.001 (Wilcoxon rank sum test). n indicates the number of embryos. Scale bars, 10 *µ*m.

**Supplementary figure 5:**
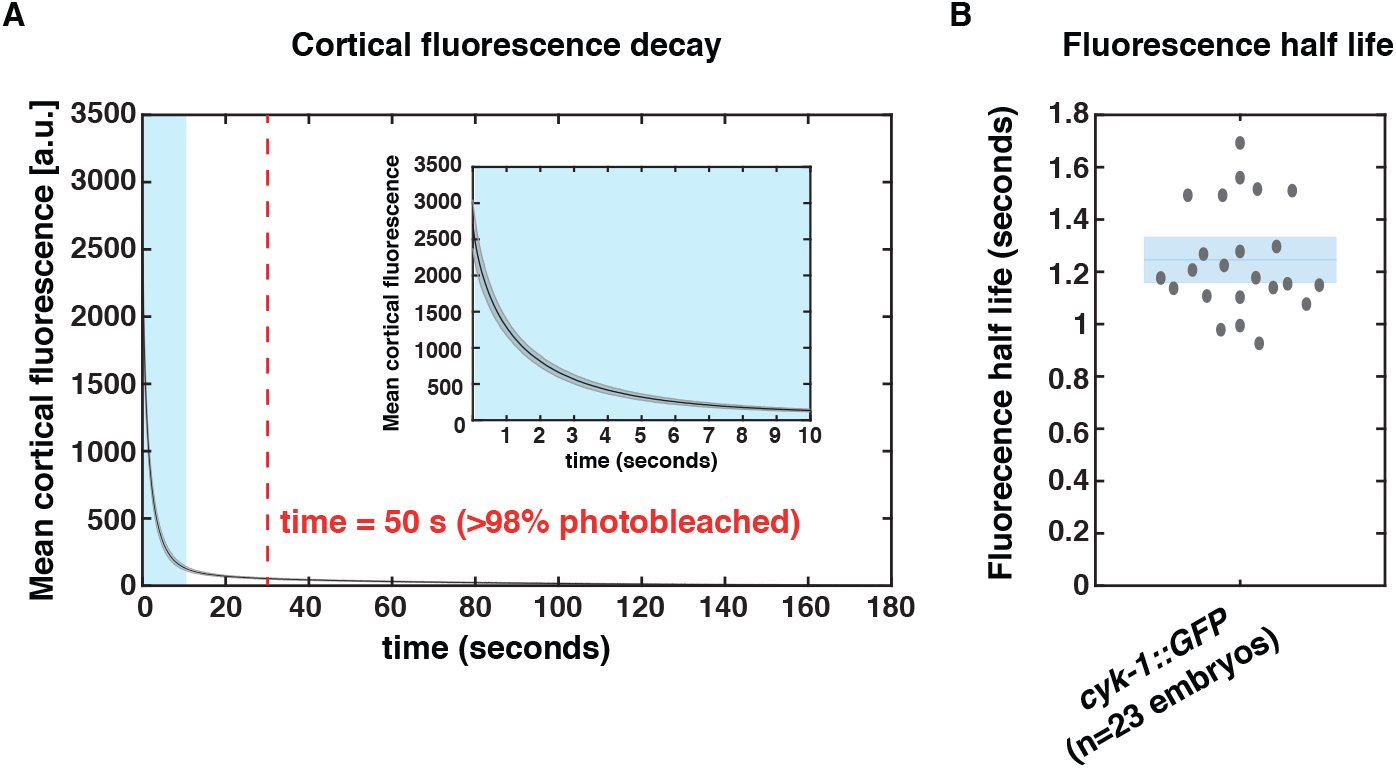
(**A**) Mean CYK-1/Formin::GFP fluorescence of 23 control embryos, measured over time during continuous spinning disc imaging. After 50 seconds of imaging 98.74 ± 0.09 % was photo bleached (indicated in red). Inset shows magnification of the light blue area in the main graph. (**B**) Fluorescence half life obtained by fitting a double exponential to fluorescence decay curves of individual embryos. Fitting yielded a fast bleaching time scale, displayed in (**B**), corresponding to the CYK-1/Formin::GFP signal, and a slow time scale corresponding to autofluorescence decay. Data points represent individual embryos. Blue box indicates the mean with 95 % confidence interval.

**Supplementary figure 6:**
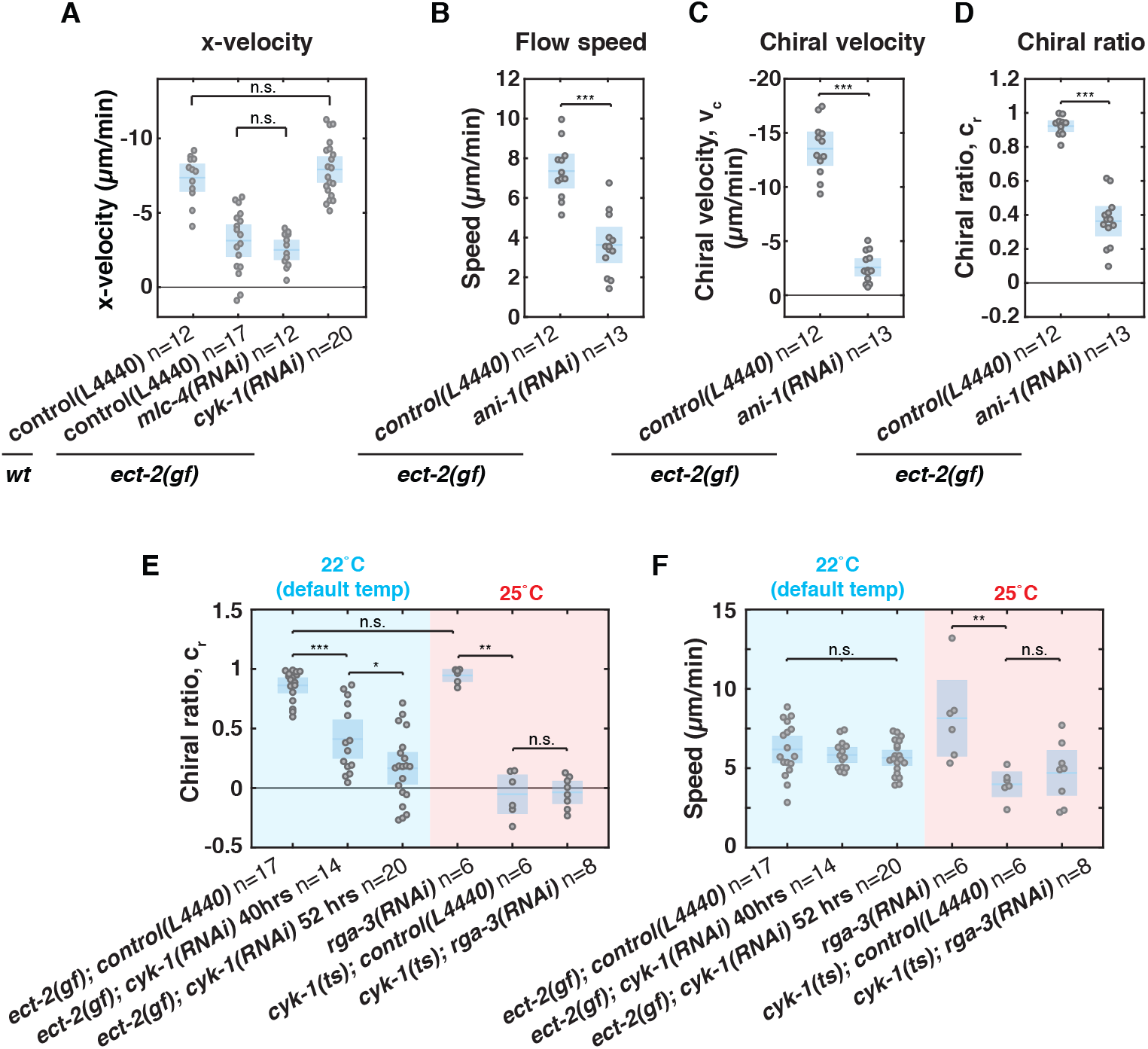
(**A**) Mean x-velocity v_x_ per embryo in wild type with control RNAi (L4440), *ect-2(gf)* with control RNAi treatment (L4440), *ect-2(gf); mlc-4(RNAi)* and *ect-2(gf); cyk-1(RNAi)*. Data points represent individual embryos. (**B**) Cortical flow speed, (**C**) chiral velocity and (**D**) chiral ratio in *ect-2(gf)* embryos on control (L4440) or *ani-1(RNAi)*. (**E**) Blue background: Chiral ratio of *ect-2(gf)* embryos on control, mild (40 hrs RNAi treatment) or strong (52 hrs RNAi treatment) *cyk-1(RNAi)* imaged at 22 °C. The ect-2(gf); control(L4440) and ect-2(gf); cyk-1(RNAi) 52 hrs data are the same as in main figure 5G. Red background: control embryos on *rga-3(RNAi), cyk-1(lf)* embryos on control and *cyk-1(lf)* embryos on *rga-3(RNAi)* imaged at 25 °C. (**F**) Blue background: Cortical flow speed of *ect-2(gf)* embryos on control, mild (40 hrs RNAi treatment) or strong (52 hrs RNAi treatment) *cyk-1(RNAi)* imaged at 22 °C. The ect-2(gf); control(L4440) and ect-2(gf); cyk-1(RNAi) 52 hrs data are the same as in main figure 5G. Red background: control embryos on *rga-3(RNAi), cyk-1(lf)* embryos on control and *cyk-1(lf)* embryos on *rga-3(RNAi)* imaged at 25 °C. In all figure panels: Mean with 95% confidence interval indicated in blue boxes. Significance testing: n.s. represents p> 0.05, * represents p *<*=0.05, ** represents p *<* =0.01, *** represents p *<* =0.001 (Wilcoxon rank sum test). n indicates number of embryos.

**Supplementary figure 7:**
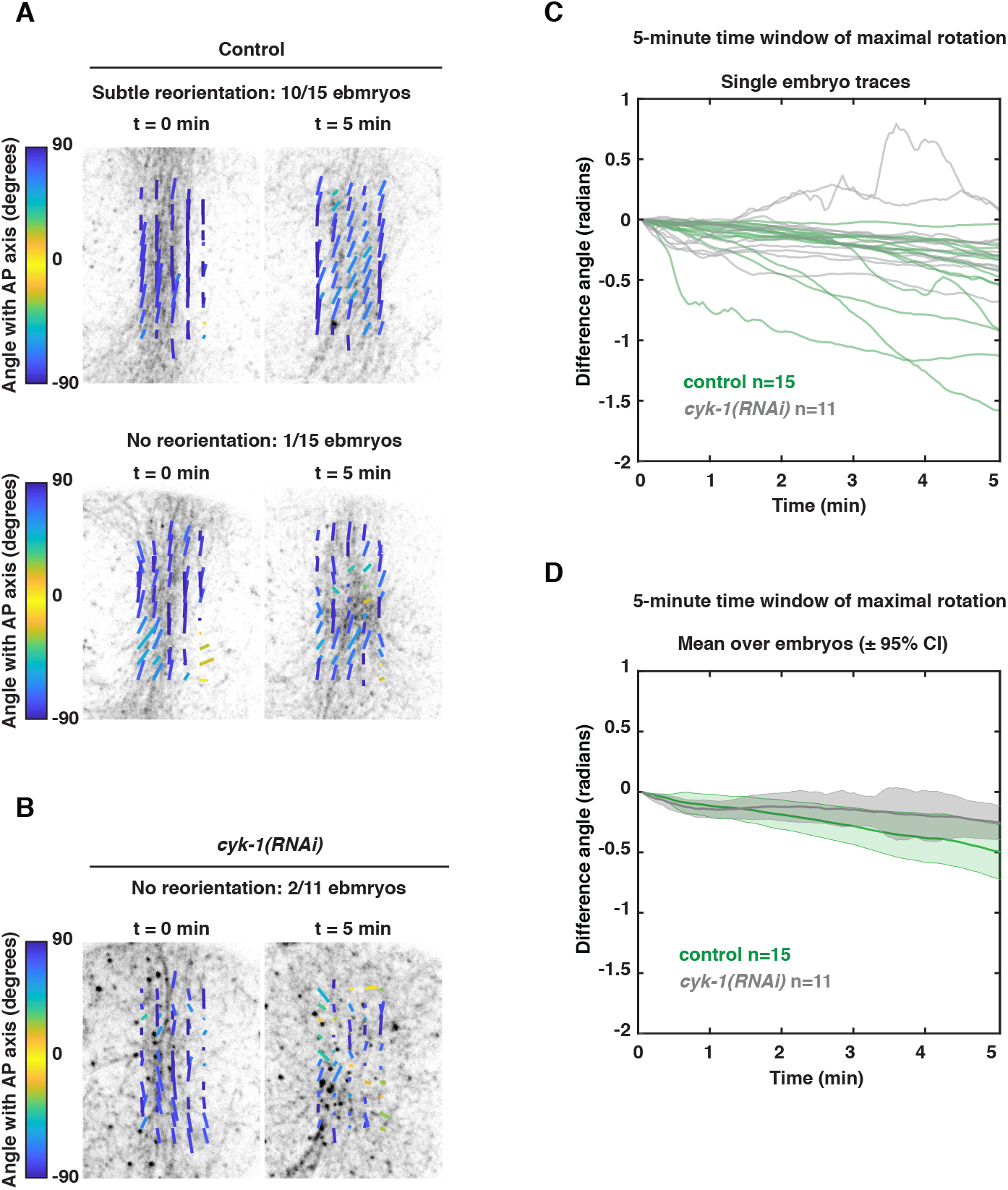
(**A-B**) Fluorescent micrographs of cortical Lifeact-mKate2 in two control embryos (**A**) and one *cyk-1(RNAi)* embryo (**B**), overlaid with the corresponding nematic field at the onset of patch formation (t=0 min) and at the end of patch rotation (t=5 min). Upper panel in (**A**) displays a mild clockwise reorientation (as observed in 10 out of 15 embryos), and the lower panel displays no reorientation (as observed in 1 out of 15 embryos). In 4 out of 15 control embryos a strong clockwise reorientation was observed (see main figure 6 for an example). The *cyk-1(RNAi)* embryo in (**B**) displays no reorientation (as observed 2 out of 11 *cyk-1(RNAi)* embryos). In 9 out of 11 *cyk-1(RNAi)* embryos a subtle clockwise rotation was observed (see main figure 6 for an example). The directors are color coded for the angle with the anteroposterior axis. (**C**) Moving average time traces of the mean actin filament orientation in the 5-minute time interval in which the clockwise reorientation was maximal. The sliding window of the moving average is 60 seconds. The vertical axis gives the difference in mean actin filament orientation with respect to the first timepoint of the 5-minute time interval. Note that, with respect to the onset of patch formation, the start of this time interval differed in each embryo (see methods). (**D**) Mean with 95% confidence interval of the traces shown in (**C**). n indicates the number of embryos.

**Supplementary figure 8:**
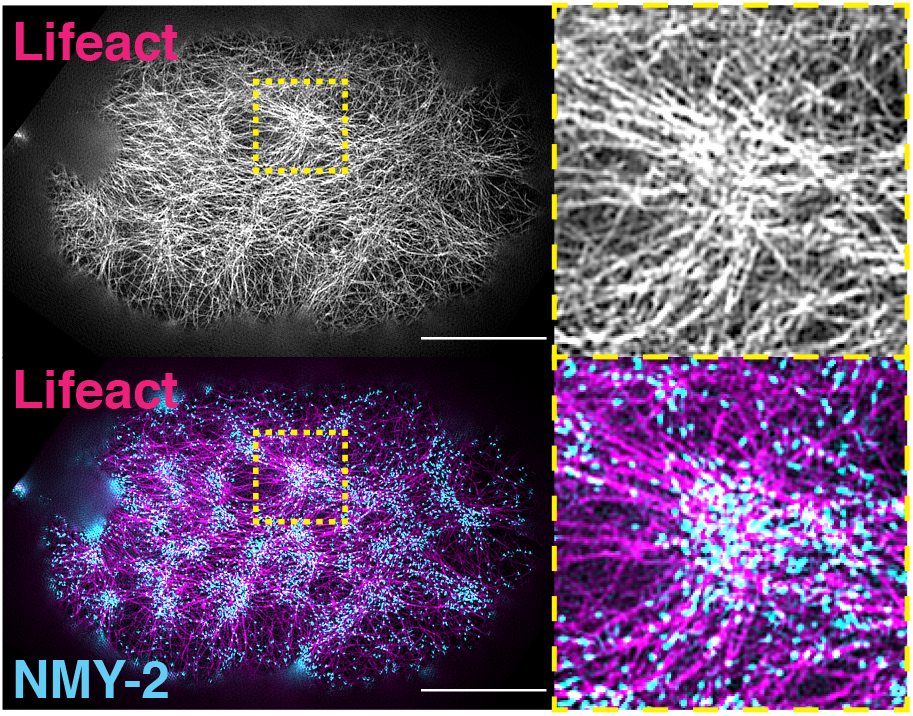
Structured Illumination Microscopy-TIRF (SIM-TIRF) micrograph of an embryo producing NMY-2::GFP and Lifeact-mKate2. Insets in the right show magnification of the regions marked with yellow boxes in left panels. Although many actin filaments are isotropically oriented, the inset shows a clear indication of an aster-like topology with an enrichment of NMY-2::GFP, as a marker for active RhoA foci, at the aster center. Scale bars, 10 *µ*m.

## 13 Movie Legends

**Movie 1 - Polarizing actomyosin flows are chiral with consistent handedness**: Cortical plane of a zygote producing endogenously labeled NMY-2 (NMY-2::GFP) imaged during polarizing flow at 25 °C with 5 second time intervals. The anterior and posterior cortical halves counter-rotate with respect to each other. Anterior is left, posterior is right in all movies, unless stated otherwise.

**Movie 2 - Actomyosin flow chirality depends on cyk-1**: Cortical plane of a *cyk-1(ts)* mutant zygote producing endogenously labeled NMY-2 (NMY-2::GFP) imaged during polarizing flow at 25 °C with 5 second time intervals. The anterior and posterior cortical halves do not counter-rotate with respect to each other.

**Movie 3 - Polarizing actomyosin flows visualized by lifeact-mKate2 upon control injection**: Cortical plane of a zygote producing Lifeact-mKate2, derived from injection of a control expression plasmid (*ph-gfp-lov2*), imaged during polarizing flow with 5 second time intervals. The anterior and posterior cortical halves counter-rotate with respect to each other.

**Movie 4 - Polarizing actomyosin flows visualized by lifeact-mKate2 in an embryo expressing high levels of CA-CYK-1/Formin** Cortical plane of a zygote producing Lifeact-mKate2, derived from injection of a *ca-cyk-1* expression plasmid, imaged during polarizing flow with 5 second time intervals. Actomyosin flow direction changes repeatedly over time.

**Movie 5 - Polarizing actomyosin flows visualized by lifeact-mKate2 in an embryo expressing low levels of CA-CYK-1/Formin** Cortical plane of a zygote producing Lifeact-mKate2, derived from injection of a *ca-cyk-1* expression plasmid, imaged during polarizing flow with 5 second time intervals. The counter-rotation of the actomyosin flow is increased when compared to embryos derived from control injections.

**Movie 6 - Left-right symmetry breaking of a 4-6 cell control embryo** Maximum intensity projection along the dorsoventral axis of a control embryo producing Tubulin::mCherry undergoing cell division of the ABa blastomere (top) and the ABp blastomere (bottom) imaged at 25 °C. Time interval is 5 seconds. Anterior is up, left side is left. Both ABa and ABp undergo a clockwise skew.

**Movie 7 - Left-right symmetry breaking is perturbed in cyk-1 mutants** Maximum intensity projection along the dorsoventral axis of a *cyk-1* mutant embryo, producing Tubulin::mCherry undergoing cell division of the ABa blastomere (top) and the ABp blastomere (bottom) imaged at 25 °C. Time interval is 5 seconds. Anterior is up, left side is left. Clockwise skew of ABa and ABp is reduced.

**Movie 8 - Polarizing flows in an embryo producing CYK-1/Formin::GFP** Cortical plane of a zygote on RNAi control (L4440) producing endogenously labeled CYK-1/Formin (CYK-1/Formin::GFP) imaged during polarizing flows with 10 second time intervals. CYK-1/Formin localizes in foci that get depleted from the posterior.

**Movie 9 - Polarizing flows in an embryo producing CYK-1/Formin::GFP upon rga-3(RNAi)** Cortical plane of a zygote on *rga-3(RNAi)* producing endogenously labeled CYK-1/Formin (CYK-1/Formin::GFP) imaged during polarizing flows with 10 second time intervals. CYK-1/Formin levels are increased with respect to controls and chiral counter-rotation of the flow is more pronounced.

**Movie 10 - Polarizing flows in an embryo producing NMY-2::GFP** Cortical plane of a zygote on RNAi control (L4440) producing endogenously labeled NMY-2 (NMY-2::GFP) imaged during polarizing flows with 5 second time intervals. NMY-2 localizes in foci, like CYK-1/Formin, that get depleted from the posterior.

**Movie 11 - Polarizing flows in an embryo producing NMY-2::GFP upon rga-3(RNAi)** Cortical plane of a zygote on *rga-3(RNAi)* producing endogenously labeled NMY-2 (NMY-2::GFP) imaged during polarizing flows with 5 second time intervals. NMY-2 levels are increased with respect to controls and chiral counter-rotation of the flow is more pronounced.

**Movie 12 - Polarizing flows in an embryo producing the active RhoA probe mCherry::ANI-1(AH-PH) and CYK-1::GFP** Cortical plane of a zygote producing mCherry::ANI-1(AH-PH) and endogenously labeled CYK-1 (CYK-1::GFP) imaged during polarizing flows with 5 second time intervals. CYK-1 and ANI-1(AH-PH) partially co-localize.

**Movie 13 - Polarizing flows in a control embryo producing NMY-2::GFP** Cortical plane of a control zygote on RNAi control (L4440) producing endogenously labeled NMY-2 (NMY-2::GFP) imaged during polarizing flows with 5 second time intervals.

**Movie 14 - Polarizing flows in an ect-2(gf) mutant embryo producing NMY-2::GFP** Cortical plane of an *ect-2(gf)* zygote on RNAi control (L4440) producing endogenously labeled NMY-2 (NMY-2::GFP) imaged during polarizing flows with 5 second time intervals. Chiral counter-rotating flows are strongly enhanced when compared to control embryos.

**Movie 15 - Polarizing flows in an ect-2(gf); mlc-4(RNAi) embryo producing NMY-2::GFP** Cortical plane of an *ect-2(gf)* zygote on *mlc-4(RNAi)* producing endogenously labeled NMY-2 (NMY-2::GFP) imaged during polarizing flows with 5 second time intervals. Cortical flow speed is reduced when compared to *ect-2(gf)* mutants on RNAi control but is still strongly counter-rotating.

**Movie 16 - Polarizing flows in an ect-2(gf); cyk-1(RNAi) embryo producing NMY-2::GFP** Cortical plane of an *ect-2(gf)* zygote on *cyk-1(RNAi)* producing endogenously labeled NMY-2 (NMY-2::GFP) imaged during polarizing flows with 5 second time intervals. *cyk-1(RNAi)* rescues the *ect-2(gf)*-induced hyperchirality phenotype.

**Movie 17 - Polarizing flows in an ect-2(gf); ani-1(RNAi) embryo producing NMY-2::GFP** Cortical plane of an *ect-2(gf)* zygote on *ani-1(RNAi)* producing endogenously labeled NMY-2 (NMY-2::GFP) imaged during polarizing flows with 5 second time intervals. *ani-1(RNAi)* affects both the overall flow speed and chirality.

**Movie 18 - Polarizing flows at 25** °**C in an rga-3(RNAi) zygote** Cortical plane of an *rga-3(RNAi)* zygote producing endogenously labeled NMY-2 (NMY-2::GFP) imaged during polarizing flows with 5 second time intervals. *rga-3(RNAi)* triggers a strong increase in actomyosin flow chirality.

**Movie 19 - Polarizing flows at 25** °**C in a cyk-1(lf) mutant** Cortical plane of a *cyk-1(lf)* mutant, on RNAi control (L4440), producing endogenously labeled NMY-2 (NMY-2::GFP) imaged during polarizing flows with 5 second time intervals.

**Movie 20 - Polarizing flows at 25** °**C upon cyk-1(lf); rga-3(RNAi)** Cortical plane of a *cyk-1(lf)* mutant on *rga-3(RNAi—)*, producing endogenously labeled NMY-2 (NMY-2::GFP) imaged during polarizing flows with 5 second time intervals. *cyk-1(lf); rga-3(RNAi)* embryos phenocopy *cyk-1(lf); control(L4440)* embryos.

**Movie 21 - Compression-induced collapse of the cytokinteic ring in an embryo producing CYK-1::GFP** Cortical plane of an embryo producing endogenously labeled CYK-1 (CYK-1::GFP) imaged during collapse of the cytokinetic ring with 3 second time intervals. CYK-1 accumulates in a patch in the center of the cortical plane.

**Movie 22 - Compression-induced collapse of the cytokinteic ring in an embryo producing NMY-2::GFP** Cortical plane of an embryo producing endogenously labeled NMY-2 (NMY-2::GFP) imaged during collapse of the cytokinetic ring with 3 second time intervals. NMY-2 accumulates in a patch the center of the cortical plane.

**Movie 23 - Compression-induced collapse of the cytokinteic ring in an embryo producing lifeact-TagRFP displaying a strong clockwise reorientation of F-actin** Cortical plane of an embryo producing lifeact-TagRFP imaged during collapse of the cytokinetic ring with 3 second time intervals. F-actin accumulates in a patch the center of the cortical plane. In addition, this embryo displays a strong clockwise reorientation of the F-actin filaments within and around the patch (as is observed in 4 out of 15 control embryos).

**Movie 24 - Compression-induced collapse of the cytokinteic ring in an embryo producing lifeact-TagRFP displaying a mild clockwise reorientation of F-actin** Cortical plane of an embryo producing lifeact-TagRFP imaged during collapse of the cytokinetic ring with 3 second time intervals. This embryo displays a mild clockwise reorientation of the F-actin filaments within and around the patch (as observed in 10 out of 15 control embryos).

**Movie 25 - Compression-induced collapse of the cytokinteic ring in an embryo producing lifeact-TagRFP displaying no observable reorientation of F-actin** Cortical plane of an embryo producing lifeact-TagRFP imaged during collapse of the cytokinetic ring with 3 second time intervals. This embryo displays no clockwise reorientation of the F-actin filaments (as observed in 1 out of 15 control embryos).

**Movie 26 - Compression-induced collapse of the cytokinteic ring in a cyk-1(RNAi) embryo producing lifeact-TagRFP displaying a mild clockwise reorientation of F-actin** Cortical plane of an embryo producing lifeact-TagRFP imaged during collapse of the cytokinetic ring with 3 second time intervals. This embryo displays a mild clockwise reorientation of the F-actin filaments (as observed in 9 out of 11 *cyk-1(RNAi)* embryos).

**Movie 27 - Compression-induced collapse of the cytokinteic ring in a cyk-1(RNAi) embryo producing lifeact-TagRFP displaying no observable reorientation of F-actin** Cortical plane of an embryo producing lifeact-TagRFP imaged during collapse of the cytokinetic ring with 3 second time intervals. This embryo displays no observable reorientation of F-actin (as observed in 2 out of 11 *cyk-1(RNAi)* embryos).

**Movie 28 - SIM-TIRF recording of cortical Lifeact-mKate2 during early polarizing flows** F-actin displays numerous inhomogeneities that resemble asters. Scale bar is 10 *µ*m.

**Movie 29 - SIM-TIRF recording of cortical Lifeact-mKate2 and NMY-2::GFP during early polarizing flows** Dual-color labeling of Lifeact-mKate2 and NMY-2::GFP. Same embryo as in Movie 28. NMY-2::GFP, localizing to regions of active RhoA, localizes at the center of aster-like structures. Scale bar is 10 *µ*m.

